# The neural code for face memory

**DOI:** 10.1101/2021.03.12.435023

**Authors:** Liang She, Marcus K. Benna, Yuelin Shi, Stefano Fusi, Doris Y. Tsao

## Abstract

The ability to recognize familiar visual objects is critical to survival. A central assumption of neuroscience is that long-term memories are represented by the same brain areas that encode sensory stimuli (*1*). Neurons in inferotemporal (IT) cortex represent the sensory percept of visual objects using a distributed axis code (*2–4*). Whether and how the same IT neural population represents the long-term memory of visual objects remains unclear. Here, we examined how familiar faces are encoded in face patch AM and perirhinal cortex. We found that familiar faces were represented in a distinct subspace from unfamiliar faces. The familiar face subspace was shifted relative to the unfamiliar face subspace at short latency and then distorted to increase neural distances between familiar faces at long latency. This distortion enabled markedly improved discrimination of familiar faces in both AM and PR. Inactivation of PR did not affect these memory traces in AM, suggesting that the memory traces arise from intrinsic recurrent processes within IT cortex or interactions with downstream regions outside the medial temporal lobe (*5, 6*). Overall, our results reveal that memories of familiar faces are represented in IT and perirhinal cortex by a distinct long-latency code that is optimized to distinguish familiar identities.

---

Our experience of the world is profoundly shaped by memory. Whether we are shopping for a list of items at the grocery store or talking to friends at a social gathering, our actions depend critically on remembering a large number of visual objects. Multiple studies have explored the molecular (*7, 8*) and cellular (*9, 10*) basis for memory, but the network-level code remains elusive. How is a familiar song, place, or face encoded by the activity of neurons?

Recent work on the sensory code for visual object identity in IT suggests that objects are encoded as points in a continuous low-dimensional object space, with single IT neurons linearly projecting objects onto specific preferred axes (*2–4*) (**Fig. 1A**, left). These axes are defined by weightings of a small set of independent parameters spanning the object space. This coding scheme (also referred to as linear mixed selectivity (*11, 12*), and related to disentangled representations in machine learning (*13*)) is efficient, allowing infinitely many different objects to be represented by a small number of neurons. Indeed, the axis code carried by macaque face patches allows detailed reconstruction of random realistic faces using activity from only a few hundred neurons (*3*).

**Figure 1.**
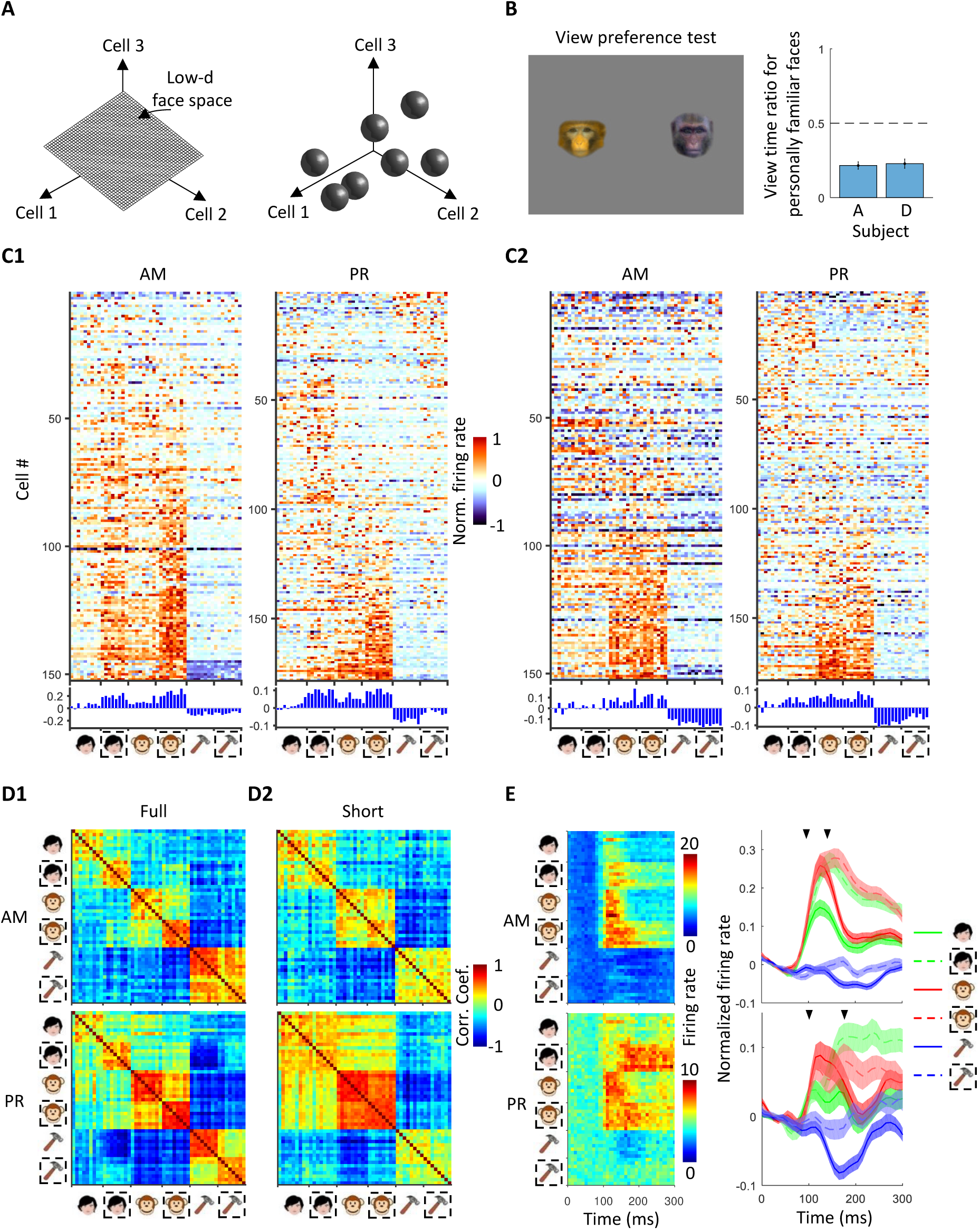
Cells in face patches AM and PR are modulated by familiarity. **(A)**Two alternative schemes for face representation: a low-dimensional continuous feature space (left) and a set of discrete attractors (right). **(B)** View preference test. Pairs of faces, one familiar and one unfamiliar, were presented for 10 s, and time spent fixating each was recorded. Right: Ratio of time spent fixating personally familiar versus unfamiliar faces for two animals. **(C1)** Responses of cells to stimuli from six stimulus categories (familiar human faces, unfamiliar human faces, familiar monkey faces, unfamiliar monkey faces, familiar objects, and unfamiliar objects) recorded from two face patches (AM, PR). Responses were averaged between 50 to 300 ms after stimulus onset (“full” response window). Dashed square, unfamiliar stimuli. P values for two-sided T-test comparing familiar vs. unfamiliar for human faces, monkey faces and objects in PR were 6.6×10^-18^, 7.7×10^-3^, 1.5×10^-7^, respectively, and in AM were 1.8×10^-23^, 2.7×10^-17^, 1.1×10^-5^, respectively. (PR: N = 1539 for familiar human/monkey faces, 1368 for unfamiliar human/ monkey faces, and 1368 for familiar/unfamiliar objects; AM: N = 1368 for familiar human/monkey faces, 1216 for unfamiliar human/monkey faces, and 1216 for familiar/unfamiliar objects.) **(C2)** Same as C1 for “short” response window (PR, 50-175 ms; AM, 50-125 ms). P values ordered as in (C1) were 0.006, 0.72, 9×10^-5^ for PR, 0.28, 0.40, 0.86 for AM. **(D1)** Similarity (Pearson correlation coefficient) matrix of population responses for full response window**. (D2)** Same as (D1) for short response window. **(E)** Left: Average response time course across AM (top) and PR (bottom) populations to each of the screening stimuli. Right: Response time course across AM and PR populations averaged across both cells and category exemplars (normalized for each cell, see Method). Earlier arrow indicates the mean time when visual responses to faces became significantly higher than baseline (AM: 92.5 ms, PR: 105 ms; see Methods). Later arrow indicates the mean time when responses to familiar versus unfamiliar faces became significantly different (AM: 125 ms and 155 ms for human faces and monkey faces, respectively; PR: 155 ms and 205 ms). Shaded areas indicate standard error of the mean (SEM) across neurons.

Here, we set out to leverage recent insight into the detailed sensory code for facial identity in IT cortex (*3*) in order to explore the population code for face memories. A longstanding assumption of neuroscience is that long-term memories are stored by the same cortical populations that encode sensory stimuli (*1*). This suggests that the same neurons that carry a continuous axis-based object coding scheme should also support tagging of a discrete set of remembered objects as familiar. However, schemes for representing discrete familiar items often invoke attractors (*14*) that would lead to breakdowns in continuous representation (**Fig. 1A**, right). This raises a key question: does familiarity alter the IT axis code for facial identity? We surmised that discovering the answer might reveal the neural code for face memory.

Previous studies have generally found decreased and sparsened responses to familiar stimuli in IT and perirhinal cortex (*15–20*). The simple view of a generalized decrease in response is at odds with the observed increase in the discriminability of familiar stimuli (*21*). A possible reason for this discrepancy is that previous recordings exploring neural correlates of visual familiarity were not targeted to specific subregions of IT cortex known to play a causal role in discrimination of the visual object class being studied (*22*).

We targeted two regions: face patch AM, the most anterior face patch in IT cortex (*23*), and face patch PR, a recently reported face patch in perirhinal cortex (*24*). These two regions lie at the apex of the macaque face patch system, an anatomically connected network of regions in the temporal lobe dedicated to face processing (*23, 25–29*). AM harbors a strong signal for invariant facial identity (*3, 23*), while perirhinal cortex plays a critical role in visual memory (*30–33*). We thus hypothesized that a representation of face memory should occur in the circuit linking AM to PR.

Our recordings revealed that familiar stimuli are encoded by a *unique geometry at long latency.* This geometry leads to increased distances between and discriminability of familiar faces, in agreement with human psychophysics (*21*). The finding that a major piece of the network code for visual memory is dynamic and only activated at long latency sheds light on how we can both veridically perceive visual stimuli and recall past experiences from them.

## Results

### AM and PR are strongly modulated by familiarity

We identified face patches AM and PR in five animals using fMRI (*25*). To characterize the role of familiarity in modulating responses of cells in AM and PR, we targeted electrodes to these two patches (**Fig. S1**) and recorded responses to a set of screening stimuli consisting of human faces, monkey faces, and objects. The stimuli were personally familiar or unfamiliar (**Fig. S2A**), with 8-9 images/category. Personally familiar images depicted people, monkeys, and objects that the animals interacted with on a daily basis; a new set of unfamiliar images was presented per cell. Animals showed highly significant preferential looking toward the unfamiliar face stimuli and away from familiar face stimuli (**Fig. 1B**), confirming behaviorally that these stimuli were indeed familiar to the monkey (*34*).

Across the population, 93% of cells in AM and 74% of cells in PR were face selective (**Fig. S3A**). Below, we group data from two monkeys for PR and three monkeys for AM, as we did not find any marked differences between individuals (**Fig. S4** shows the main results separately for each animal). Both AM and PR exhibited a significantly stronger response across the population to unfamiliar compared to personally familiar stimuli in this experiment (**Fig. 1C1**; full response distributions are shown in **Fig. S3B**). This is consistent with a large number of previous studies reporting suppression of responses to familiar stimuli in IT and perirhinal cortex (*15–20*) (though it is discrepant with a recent monkey fMRI study reporting a stronger response to familiar compared to unfamiliar faces in all temporal lobe face patches (*24*)). Individual cells showed a diversity of selectivity profiles for face species and familiarity type (**Fig. S5A, B**). The local field potential in both areas was also face-selective and significantly stronger to unfamiliar compared to personally familiar faces (**Fig. S5C**). Representation similarity matrices revealed distinct population representations of the six stimulus classes in both AM and PR (**Fig. 1D1**); this was confirmed by multidimensional scaling analysis (**Fig. S6**).

Mean responses to familiar versus unfamiliar faces diverged over time, with the difference becoming significant at 125 ms in AM and 175 ms in PR; the mean visual response to faces themselves significantly exceeded baseline earlier, at 85 ms in AM and 100 ms in PR (**Fig. 1E**, **S7**). The delay in suppression to familiar faces is consistent with previous reports of delayed suppression to familiar stimuli in IT (*15, 17–19*). Interestingly, PR (and to a lesser extent, AM) responses were suppressed not only by familiar faces but also familiar objects, suggesting a role for these areas in non-face coding, possibly through associative memory mechanisms (*30*). Single-cell response profiles and representation similarity matrices computed using a short time window (50 - 125 ms after stimulus onset for AM and 50 - 175 ms for PR) showed less distinct responses to familiar versus unfamiliar stimuli (**Fig. 1C2, D2, S6**). Overall, the results so far show that both AM and PR exhibit long latency suppression to familiar faces.

**Figure 2.**
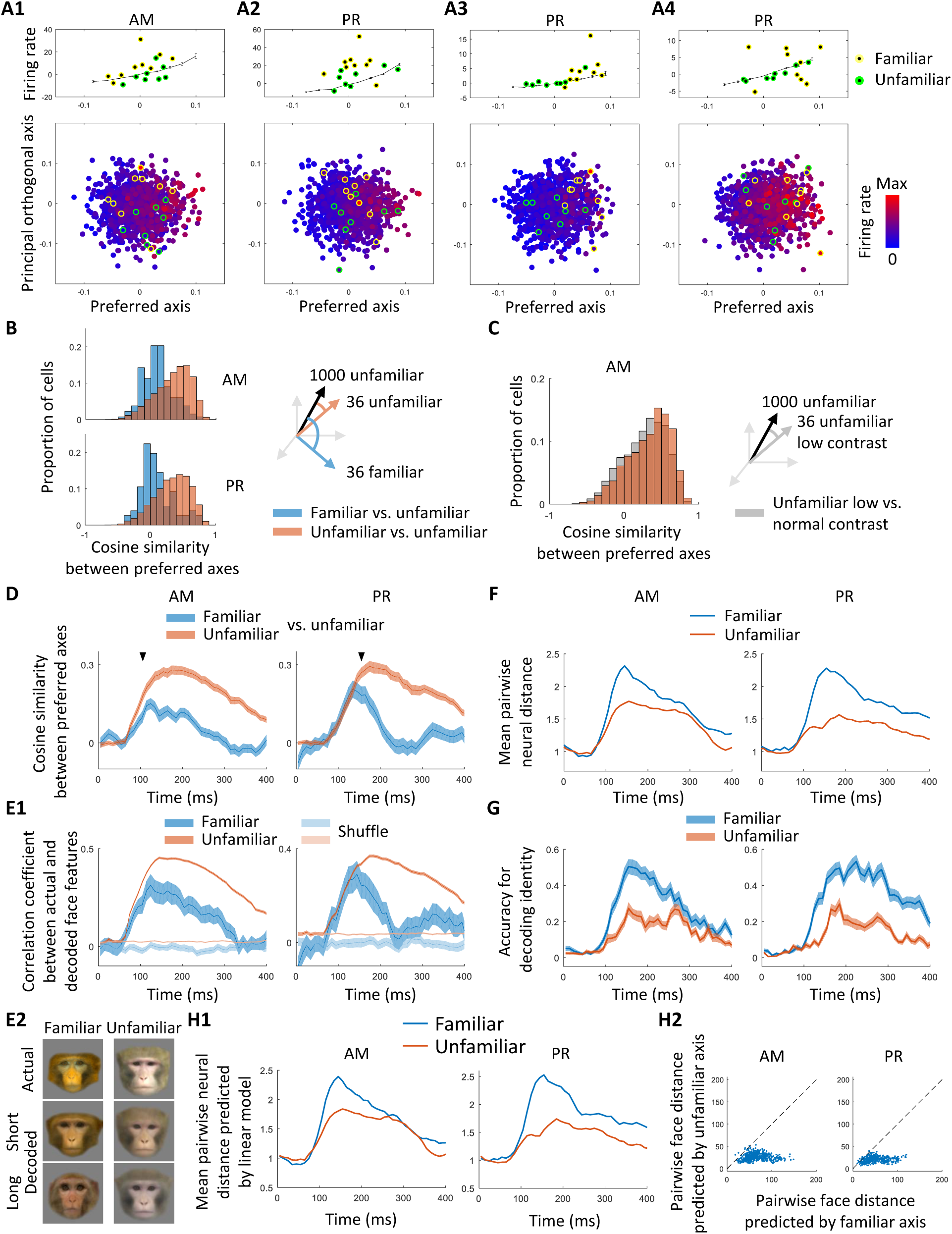
AM and PR cells use different axes to represent familiar versus unfamiliar faces. **(A)** Four example cells showing axis tuning. Bottom: Responses to 1000 unfamiliar faces, projected onto the cell’s preferred axis and principal (longest) orthogonal axis in the face feature space. Response magnitudes are color-coded. Top: Mean response as a function of distance along the preferred axis. Green (yellow) dots: projections of responses to 8 random unfamiliar (9 personally familiar) faces. Note that the 8 random unfamiliar faces indicated by the green dots were excluded from calculation of the preferred axis of the cells here. **(A1)** Axis-tuned cell from AM. **(A2-A4)** Axis-tuned cells from PR. **(B)** Population analysis comparing preferred axes for familiar versus unfamiliar faces. Top: Distribution of cosine similarities between axes computed using 1000-36 unfamiliar faces and 36 left out unfamiliar faces (orange), and between axes computed using 1000-36 unfamiliar faces and 36 familiar faces (blue). Results from 100 repeats of random subsets of 36 unfamiliar faces are averaged. Preferred axes were computed using the top 10 shape and top 10 appearance features of presented faces. **(C)** Control experiment repeating the analysis of (B) with 36 low contrast faces instead of 36 familiar faces. **(D)** Time course of the cosine similarity between preferred axes for unfamiliar-unfamiliar (orange) and unfamiliar-familiar (blue) faces; same as (B), but computed using a 50 ms sliding time window, step size 10 ms. Arrows indicate when differences became significant (AM: 105 ms, PR: 155 ms, one-tailed T-test, p<0.001, AM N=134, PR N=76). Shaded area, SEM. **(E1)** Time course of linear decoding performance for familiar and unfamiliar faces measured by the correlation coefficient between actual and decoded face feature vectors, computed using a 50 ms sliding time window, step size 10 ms. Shaded area, SEM. Light color, same analysis using stimulus identity-shuffled data (10 repeats). **(E2)** Example linearly reconstructed faces from short (120-170 ms) or long (220-270 ms) latency responses combining cells from both PR and AM. Reconstructions were performed using decoders trained on unfamiliar faces. **(F)** Time course of mean pairwise neural distance (Euclidean distance between population responses) between familiar or unfamiliar faces, computed using a 50 ms sliding time window, step size 10 ms, normalized by mean baseline (0-50 ms) distance between unfamiliar faces. Distances were computed using a subset of 30 familiar and unfamiliar feature-matched faces (see **Fig. S14**). **(G)** Time course of face identity decoding accuracy for 30 familiar (blue) or unfamiliar (orange) feature-matched faces, computed using a 50 ms sliding time window, step size 10 ms. Half the trials were used to train a linear classifier and decoding performance was tested on the remaining half of trials; chance performance was 1/30. The difference between familiar and unfamiliar identity decoding arose before the long-latency axis change in PR (115 ms in both AM and PR, one-tail paired T-test, p<0.01), possibly due to higher response reliability for familiar faces (**Fig. S16B**). **(H1)** Time course of mean pairwise neural distance between 30 familiar or unfamiliar feature-matched faces, normalized by mean baseline (0-50 ms) distance between unfamiliar faces, computed using responses predicted by linear model. Same conventions as (F). **(H2)** Mean pairwise distances between 30 familiar faces predicted using familiar axes (x-axis) and unfamiliar axes (y-axis) (see Methods).

### AM and PR use an axis code to represent unfamiliar facial identity

Responses of AM and PR cells to familiar stimuli, while lower on average at long latencies, remained highly heterogeneous across faces (**Fig. 1C1**, **S5A, B**), indicating that they were driven by both familiarity and identity. We next asked how familiarity interacts with the recently discovered axis code for facial identity.

According to the axis code, face cells in IT compute a linear projection of incoming faces formatted in shape and appearance coordinates onto specific preferred axes (*3*) (**Fig. S8A**); for each cell, the preferred axis is given by the coefficients 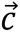 in the equation 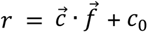 where *r* is the response of the cell, 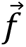 is a vector of shape and appearance features, and *c_0_* is a constant offset (see Methods). Together, a population of face cells with different preferred axes encodes a face space that is embedded as a linear subspace of the neural state space (**Fig. S8A**). The axis code has so far been examined only for unfamiliar faces. By studying whether and how this code is modified by familiarity, we reasoned that we could potentially understand the code for face memory.

We first asked whether face cells encode familiar and unfamiliar faces using the same axis. To address this, we examined tuning to unfamiliar faces (described in this section) and then compared to tuning to familiar faces (described in the next section). We began by mapping preferred axes of AM and PR cells using a set of 1000 unfamiliar monkey faces (**Fig. S2B**). We used monkey faces because responses to the screening stimuli were stronger to monkey than human faces on average in AM/PR (**Fig. 1C1,** p < 0.002, T-test, N = 323 cells pooled across AM and PR). The 1000 monkey faces were randomly drawn from a monkey face space defined by 120 parameters (see Methods), and encompassed a large variety of identities, allowing a subset to be chosen that were matched in feature distributions to familiar faces (see **Fig. S14** below).

As expected, cells in AM showed ramp-shaped tuning along their preferred axes and flat tuning along orthogonal axes (**Fig. 2A1**, **S9A, G**; additional example cells are shown in **Fig. S10A**; a control experiment with slower timing and dense sampling across two dimensions confirmed ramp-shaped tuning, **Fig. S11** and Supplementary Text). Interestingly, a large proportion of cells in PR also showed ramp-shaped tuning along their preferred axes and flat tuning along orthogonal axes (**Fig. 2A2-A4**, **S9A, G**; additional examples are shown in **Fig. S10B**). To our knowledge, this is the first time axis coding of visual features has been reported for perirhinal cortex. In both AM and PR, preferred axes computed using split halves of the data were highly consistent (**Fig. S9B**). Face identification based on feature reconstruction using linear regression revealed high performance in PR, similar to that in AM (**Fig. S9H**). These results suggest that AM and PR share a common axis code for representing unfamiliar faces.

In addition to cells with axis tuning, we found a small population of cells that responded extremely sparsely to just a few of the stimuli (**Fig. S12**, Supplementary Text). These cells constituted a very small proportion of cells in both AM (5/187) and PR (11/137). In contrast, a large proportion of cells in both areas showed significant axis tuning (108/134 cells in AM and 62/72 cells in PR, see Methods).

### Familiarity causes AM and PR responses to go off axis

We next examined how familiarity modulates the axis code. We projected the features of personally familiar and a random subset of unfamiliar faces onto the preferred axis of each AM/PR cell and plotted responses. Strikingly, responses to unfamiliar faces followed the axis (**Fig. 2A**, **S10**, green dots), whereas responses to familiar faces departed from the axis (**Fig. 2A**, **S10**, yellow dots).

This departure was not a simple gain change: the strongest responses to familiar faces were often to faces projecting somewhere in the middle of the ramp rather than on the end (**Fig. 2A**). Thus this departure cannot be explained by an attentional increase or decrease to familiar faces, which would elicit a gain change (*35*). Indeed, the effect cannot be explained by any monotonic transform in response, such as repetition suppression or sparsening (*18, 20*), as any such transform should preserve the rank ordering of preferred stimuli (**Fig. S8B**).

The surprising finding of off-axis responses to familiar faces was prevalent across the AM and PR populations. To quantify this phenomenon at the population level, we first created a larger set of familiar faces. To this end, animals were shown face images and movies daily for at least one month, resulting in a total of 36 familiar monkey faces, augmenting the 9 personally familiar monkey faces in our initial screening set (**Fig. S2C**; see Methods). Preferential looking tests confirmed that the pictorially and cinematically familiar faces were treated similarly to the personally familiar faces (**Fig. S13**). These 36 familiar faces were presented randomly interleaved with the 1000 unfamiliar monkey faces while we recorded from AM and PR.

We computed preferred axes for cells using responses to the 36 familiar faces. We found that these familiar axes significantly explained responses to familiar faces for a subset of cells (**Fig. S9C, D**). Because we only had 36 familiar faces, the statistical power for computing the preferred axis was much less than we had with 1000 unfamiliar faces, leaving open the possibility that a significant part of the familiar response may be driven by nonlinear interactions between features rather than linear axis tuning. Nevertheless, when familiar and unfamiliar face numbers were matched, familiar axes did as well as unfamiliar axes in explaining responses to faces (**Fig. S9C**). The comparable strength of axis tuning for familiar and unfamiliar faces naturally raised the question: are familiar and unfamiliar axes the same?

To compare familiar and unfamiliar axes, for each cell, we first computed the preferred axis using responses to the large set of unfamiliar parameterized faces (1000-36 faces). We then correlated this to a preferred axis computed using responses to (i) the set of 36 familiar faces (‘unfamiliar-familiar’ condition) or (ii) the left out set of 36 unfamiliar faces (‘unfamiliar-unfamiliar’ condition). The distribution of correlation coefficients revealed significantly higher similarities for the unfamiliar-unfamiliar compared to the unfamiliar-familiar condition (**Fig. 2B**).

As a control, we presented a set of low-contrast faces expected to elicit a simple decrease in response gain while preserving rank ordering of preferred stimuli. Confirming expectation, axis similarities computed using these contrast-varied faces were not significantly different for high-high versus high-low contrast faces (**Fig. 2C**). As a second control, to ensure that the effects were not due to differences in the feature content of familiar versus unfamiliar faces, we identified 30 familiar and 30 unfamiliar faces that were precisely feature-matched (see Methods and **Fig. S14**). We recomputed unfamiliar-familiar and unfamiliar-unfamiliar correlations and continued to find that familiar faces were encoded by a different axis than unfamiliar faces (**Fig. S15A**).

Earlier, we had observed that the decrease in firing rate for familiar faces occurred at long latency (**Fig. 1E**). We next investigated the time course of the deviation in preferred axis. We performed a time-resolved version of the analysis in **Fig. 2B**, comparing the preferred axis computed from 36 unfamiliar or 36 familiar faces with that computed from 1000-36 unfamiliar faces over a rolling time window (**Fig. 2D**). Initially, axes for familiar and unfamiliar faces were similar. But at longer latency (t > 105 ms in AM, t > 155 ms in PR), the preferred axis for familiar faces diverged from that for unfamiliar faces.

The divergence in preferred axis over time for familiar versus unfamiliar faces suggests that the brain would need to use a different decoder for familiar versus unfamiliar faces at long latencies. Supporting this, in both AM and PR, at short latencies, feature values for familiar faces obtained using a decoder trained on unfamiliar faces matched actual feature values, and reconstructions were good (**Fig. 2E**). In contrast, a decoder trained on unfamiliar faces at long latency performed poorly on recovering feature values of familiar faces (**Fig. 2E**; note, however, training a decoder using both familiar and unfamiliar faces at long latency yielded above-chance feature decoding accuracy for both, indicating that preferred axes for familiar and unfamiliar faces were not completely orthogonal in all cells at long latency, **Fig. S16A**).

### Axis change increases the neural distance between familiar faces

What computational purpose could the deviation in preferred axis for familiar versus unfamiliar faces serve? We hypothesized that this might allow familiar faces to be better distinguished from each other by increasing the distance between them in the neural state space. In an extreme case, imagine a pair of twins differing in only a single face feature: If preferred axes of cells weighed each feature equally, response vectors to the two faces would be almost identical, while aligning cells’ preferred axes to the distinguishing feature axis would increase the distance.

Supporting the hypothesis that axis change serves to increase discriminability of familiar faces, mean pairwise distances between neural responses to faces were indeed higher for familiar compared to unfamiliar face pairs, and the difference was greater at long latency (**Fig. 2F**). Consistent with this, identity decoding for familiar faces using a multi-class SVM was significantly better than that for unfamiliar faces (**Fig. 2G**). A linear axis model accounting for axis direction and gain changes could explain most of this increase in neural distance for familiar faces (**Fig. 2H1, Fig. S17**) (note, however, nonlinear components could still contribute to differences between familiar and unfamiliar responses, since such differences might be averaged out when computing mean pairwise distance).

Is it essential to account for axis change (**Fig. 2B, D**) in order to explain the increase in neural distance between familiar faces (**Fig. 2F**)? To address this, for each pair of familiar faces, we predicted the pairwise distance using (i) the familiar axis, or (ii) the unfamiliar axis of each cell, computed using the long-latency response (150-300 ms). Predicted pairwise distances were significantly greater for (i) compared to (ii) (**Fig. 2H2)**; if cells showed no axis change, then pairwise distances for (i) and (ii) should be the same. This confirms that the preferred axes of cells are different for familiar versus unfamiliar faces, and accounting for this difference is essential to explain the increase in neural distance between familiar faces.

### An early shift in coding subspace for familiar versus unfamiliar faces

So far, we have uncovered a new geometric mechanism to improve discrimination between familiar faces. But how is familiarity itself encoded in AM and PR? Previous studies suggest that familiarity is encoded by response suppression across cells (*15–20*). Supporting this, our first experiment revealed a decreased average response to familiar compared to unfamiliar faces (**Fig. 1**). However, to our great surprise, data from our second experiment (**Fig. 2**) showed a *stronger* mean response to familiar compared to unfamiliar stimuli (**Fig. 3A, B**). This was true even when we compared responses to the *exact same subset of images* (**Fig. S18A**). What could explain this reversal? The two experiments had one major difference: in the first experiment, the ratio of familiar to unfamiliar faces was 34/16, while in the second experiment the ratio was 36/1000 (in both experiments, stimuli were randomly interleaved and presentation times were identical). This suggests that mean response magnitude is not a robust indicator of familiarity, as it depends on temporal context. Even more challenging to the repetition suppression model of familiarity coding, the accuracy for decoding familiarity rose above chance extremely early, starting at 95 ms in AM and 105 ms in PR (**Fig. 3C**), *before* any significant difference in mean firing rates between familiar and unfamiliar faces had even emerged in either AM or PR (compare black arrow in **Fig. 3C** with green arrow in **Fig. 3B**).

**Figure 3.**
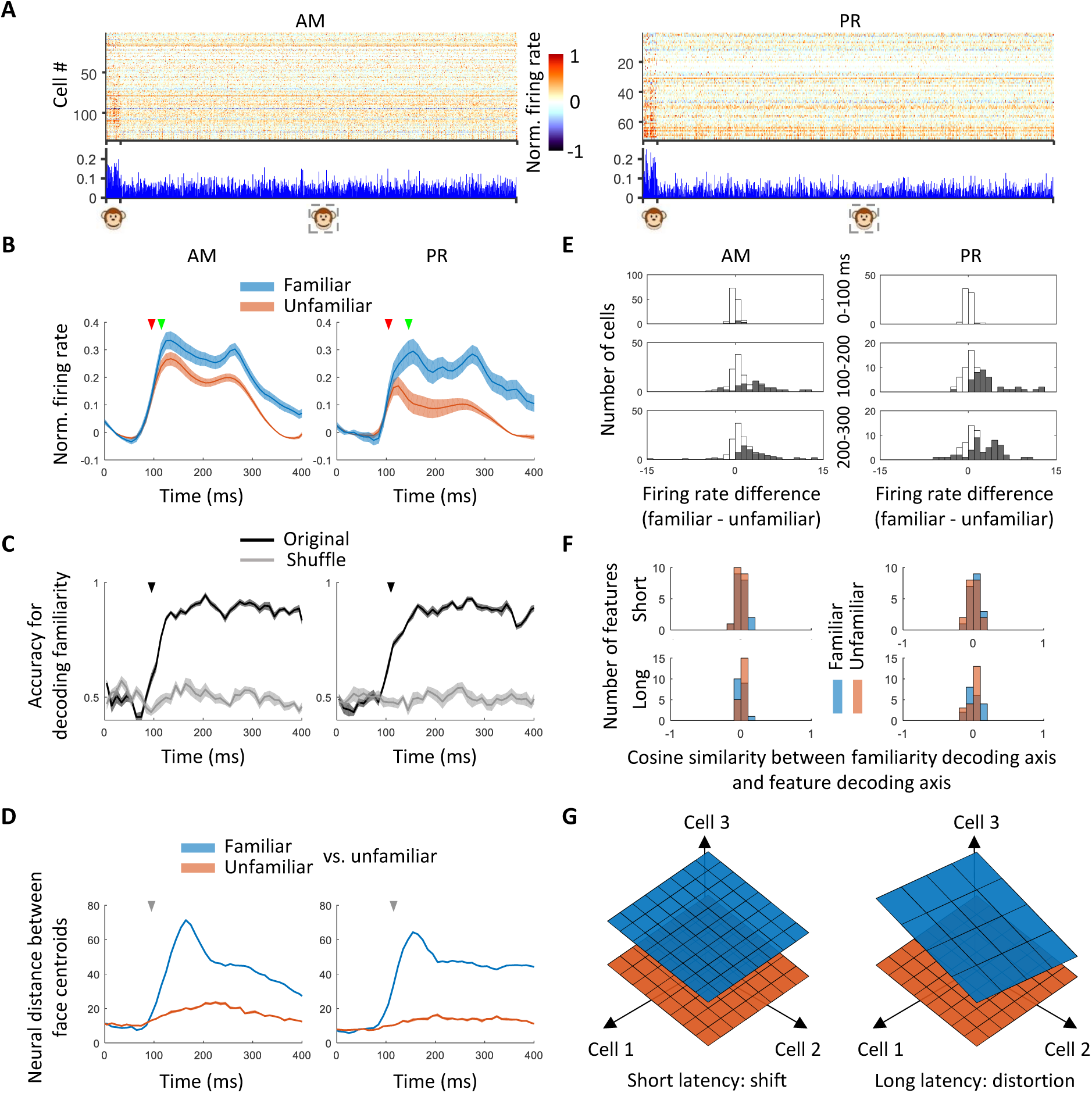
An early shift in response subspace allows familiarity to be decoded. **(A)**Responses of cells to stimuli from 36 familiar and 1000 unfamiliar monkey faces, averaged between 50 to 300 ms after stimulus onset. Dashed square, unfamiliar stimuli. **(B)** Response time course across AM and PR populations, averaged across cells and all familiar or unfamiliar faces from the 1036 monkey face stimulus set (normalized for each cell, see Methods). Shaded area, SEM. Earlier arrow indicates the time when visual responses to faces became significantly higher than baseline (AM 95 ms, PR 105 ms, one-tailed paired T-test, p<0.001, AM N=134, PR N=76). Later arrow indicates the time when responses to familiar versus unfamiliar faces became significantly different (AM 115 ms, PR 145 ms, two-sided paired T-test, p<0.001, AM N=134, PR N=76). **(C)** Time course of accuracy for decoding familiarity (black line) computed using a 50 ms sliding time window, step size 10 ms. Shaded area, SEM. Chance level was obtained using shuffled data (gray line). Arrow indicates the time at which decoding accuracy rose above chance (95 ms for AM, 105 ms for PR, one-tailed T-test, p<0.01, N=10). **(D)** Time course of neural distance between centroids of 36 familiar and 1000 unfamiliar face responses (blue) and between centroids of responses to a subset of 36 unfamiliar faces and responses to remaining 1000-36 unfamiliar faces (orange), computed using a 50 ms sliding time window, step size 10 ms. Arrow indicates the time when *d’* along the two centroids became significantly higher than a shuffle control (AM 95 ms, PR 115 ms, one-tailed T-test, p<0.01, N=10, see Methods). **(E)** Distribution of differences between mean firing rates to familiar and unfamiliar faces at 3 different time intervals. Gray bars indicate cells showing a significant difference (two-sided T-test p<0.05, familiar N=36, unfamiliar N=1000). **(F)** Distribution of cosine similarities between the familiarity decoding axis and face feature decoding axes (familiar: blue, unfamiliar: orange) at short (50-150 ms) and long (150-300 ms) latency for the first 20 features (10 shape, 10 appearance). **(G)** Schematic illustration of neural representation of familiar (blue) and unfamiliar (orange) faces at short and long latency.

What signal could support this ultra-fast decoding of familiarity, if not mean firing rate difference? Recall earlier, we had found that at short latency, familiar faces were encoded using the same axes as unfamiliar faces (**Fig. 2D**), and familiar face features could be readily decoded using a decoder trained on unfamiliar faces (**Fig. 2E**). This means that familiar and unfamiliar faces are represented in either identical or parallel manifolds at short latency. This suggested to us that their representations might be *shifted* relative to each other, and this shift is what permits early familiarity decoding. A plot of the neural distance between familiar and unfamiliar response centroids over time supported this hypothesis (**Fig. 3D**): the familiar-unfamiliar centroid distance increased extremely rapidly compared to the unfamiliar-unfamiliar one, and the *d*′ along the unfamiliar-familiar centroid axis became significantly higher than a shuffle control at 95 ms in AM and 115 ms in PR, comparable to the time when familiarity could be decoded significantly above chance. Direct inspection of shifts between responses to familiar versus unfamiliar faces across cells revealed a distribution of positive and negative values which could be exploited by a decoder for familiarity (**Fig. 3E**).

Further supporting the shift hypothesis, we found that the familiarity decoding axis was orthogonal to the face feature space at both short and long latency. We computed the cosine similarity in the neural state space between the familiarity decoding axis and face feature decoding axes, both familiar and unfamiliar, for 20 features capturing the most variance. The resulting values were tightly distributed around 0 at both short (50-150 ms) and long (150-300 ms) latency, indicating that the shift vector was orthogonal to the feature coding subspace at both latencies (**Fig. 3F**). The orthogonality between the familiarity decoding axis and the face feature space argues against familiarity being analogous to a contrast/repetition suppression signal that simply changes the gain of responses (*20*), since such a signal would lie within the feature space (note, however, a gain change that changes both the gain and offset of responses is compatible with familiarity-gated feature space shift). Overall, these results suggest a geometric picture in which familiar and unfamiliar stimuli are represented in distinct subspaces, with the familiar face subspace shifted relative to the unfamiliar face subspace at short latencies and then further distorted at long latencies to increase the distance between distinct familiar faces (**Fig. 3G**).

### Localizing the site of face memory within the face patch network

Is the distinct representation of familiar faces at long latency in AM due to feedback from PR? To address this, we silenced PR while recording responses to familiar and unfamiliar faces in AM (**Fig. 4A**). IT cortex is known to receive strong feedback from perirhinal cortex (*36*), and this is true in particular for face patch AM (*29*). Consistent with this, inactivation of PR produced strong changes in AM responses, with some cells showing an increase in response and others showing a decrease (**Fig. 4B, C, S19A**).

**Figure 4.**
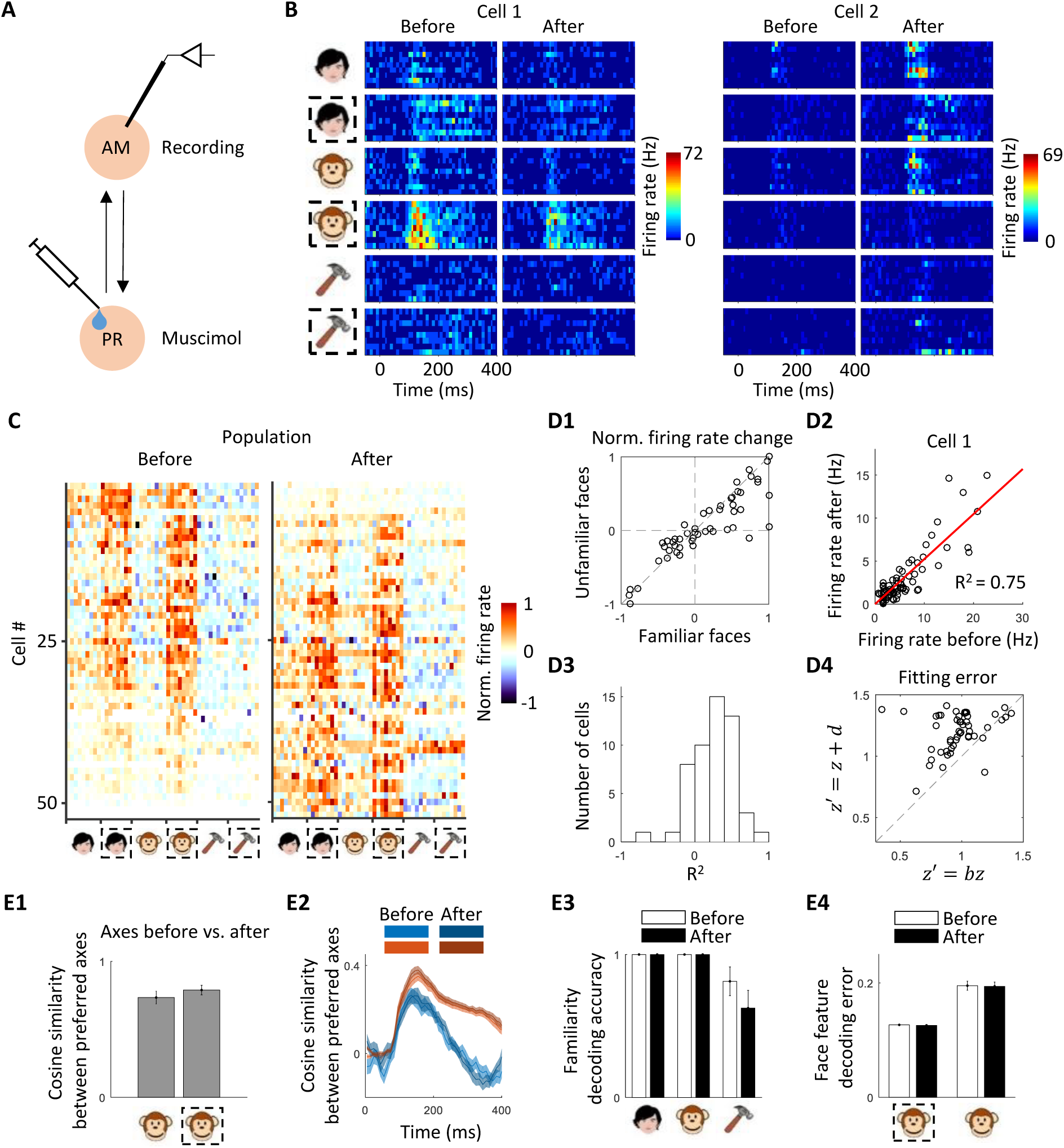
Identifying the source of the memory signal. **(A)**Schematic of experiment to identify the origin of memory-related signals in AM. Muscimol was injected into face patch PR and responses of AM neurons were recorded both before and after PR inactivation using a multi-electrode probe. **(B)** Responses of two example AM cells to screening stimuli before and after PR inactivation. Icon conventions as in Fig. 1. **(C)** Response profiles of the AM population to the screening stimuli before and after PR inactivation. **(D1)** Normalized firing rate changes for familiar and unfamiliar faces induced by PR inactivation (N = 52 cells). **(D2)** Responses of one example cell in (B) after versus before PR inactivation, fit to a linear gain function (*y = ax*). **(D3)** Distribution of *R^2^* values from fitting a linear gain function for all cells. **(D4)** Responses of each cell to the 1000 face set were fit using a gain model and an offset model (N = 50 cells). Residual errors for the offset model (ordinate) are plotted against those of the gain model (abscissa). **(E1)** Mean cosine similarity between preferred axes before and after PR inactivation for familiar or unfamiliar monkey faces. **(E2)** Time course of the similarity between preferred axes for unfamiliar-unfamiliar and unfamiliar-familiar faces (same as Fig. 2D) before and after PR inactivation. **(E3)** Familiarity decoding accuracy for screening stimuli before and after PR inactivation, using response time window 50-300 ms. **(E4)** Face feature decoding error (mean square error) for unfamiliar and familiar faces before and after PR inactivation, using response time window 50-300 ms.

We next asked whether feedback modulation from PR specifically affects responses to familiar faces, as one might expect if PR were the source of AM memory signals. We found, to the contrary, that responses to familiar and unfamiliar faces were similarly modulated by PR inactivation across the population (**Fig. 4D1**). The effect of PR inactivation was well modeled by gain change (**Fig. 4D2-D4**). Preferred axes computed using responses to 1036 monkey faces were highly similar before and after PR inactivation, for both unfamiliar and familiar faces (**Fig. 4E1**). The time course for axis divergence for familiar versus unfamiliar faces was also highly similar before and after PR inactivation (**Fig. 4E2**). Finally, decoding of both face familiarity and face features from AM activity was unaffected by PR inactivation (**Fig. 4E3**, **E4**). Overall, these results show that inactivation of PR has a strong effect on the gain of AM responses, but no apparent effect on face coding (see Supplementary Text and **Fig. S19B-D** for an interpretation of the gain changes within a hierarchical generative inference framework). In particular, the signatures of face memory we identified, namely, centroid shift and axis change, do not appear to depend on feedback from perirhinal cortex. They must thus either be intrinsic to IT cortex or arise from interactions with downstream regions outside the medial temporal lobe.

Do these signatures of familiarity exist even earlier in the face patch pathway? We mapped responses to familiar and unfamiliar faces in face patch ML, a hierarchically earlier patch in the macaque face processing pathway that provides direct input to AM (*23, 29*). Responses to the screening stimuli in ML exhibited a similar pattern as in AM, showing suppression to personally familiar faces at long latency (**Fig. S20A, C**). However, population representation similarity matrices did not show distinct population responses to familiar versus unfamiliar faces (**Fig. S20B**). Furthermore, the population average firing rate showed a sustained divergence much later than in AM (160 ms compared to 140 ms in AM, **Fig. S20C**), suggesting ML may receive a familiarity-specific feedback signal from AM. Overall, representations of familiar versus unfamiliar faces were much more similar in ML compared to AM (**Fig. S20D-K**). These results suggest ML plays a smaller role than AM and PR in storing memories of faces.

## Discussion

In this paper, we investigated the neural code for face memory in face patch AM of IT cortex and face patch PR of perirhinal cortex. PR, like AM, contained a high concentration of face-selective cells. In both regions, the vast majority of cells showed axis tuning to unfamiliar faces. At short latency, responses to familiar faces were well predicted by axes computed using unfamiliar faces, modulo a shift in the overall response subspace (**Fig. 2E**); this subspace shift enabled early detection of familiarity (**Fig. 3C-E**). We observed a striking change in the preferred axis of cells at long latency for familiar versus unfamiliar faces (**Fig. 2B, D**). The axes of individual cells changed to produce a large increase in neural population distance between distinct familiar faces (**Fig. 2F-H**). Inactivation of PR did not affect these memory-related dynamics in AM (**Fig. 4**). A simple computational model could recapitulate all of our main findings (**Fig. S21**, Supplementary Text).

These results provide the first detailed geometric picture of how visual memories are encoded. They supply a neurophysiological correlate for the behavioral finding that familiar faces can be identified much more readily than unfamiliar ones (*21*), namely, increase in neural distance between familiar faces.

Our results challenge two previous models for encoding of visual familiarity. First, contrary to the repetition suppression model for familiarity encoding (*15–20*), we found that mean response amplitude was not a robust indicator of familiarity in either patch, as it was highly sensitive to temporal context (compare **Fig. 1C1** vs. **Fig. 3A, Fig. 1E** vs. **Fig. 3B**), and emerged only after familiarity could already be decoded (**Fig. 3C**). We also note that the repetition suppression model only addresses encoding of familiarity (one bit of information) and does not address encoding of features of familiar objects, a central focus of the present study. Second, our findings challenge the sparsification model for encoding of familiar objects at long latency, which posits that neurons encode only a small number of maximally effective familiar stimuli compared to unfamiliar stimuli (*18*). We found that familiar faces triggered axis change, a fundamentally different transformation from sparsification that scrambles the rank ordering of preferred stimuli compared to that predicted by the unfamiliar axis. Overall, we believe we have identified a fundamentally new mechanism for encoding visual memory. While so far we have only characterized this mechanism within face patches, the similarity of functional organization and coding principles between face patches and other parts of IT (*4, 37*) suggests the mechanism is likely to generalize.

Face patch AM lies at the apex of the ventral form representation pathway (*23*), while perirhinal cortex, in which face patch PR is embedded, contributes to both perceptual and mnemonic functions, with a special role in visual associative memory (*30–33, 38*). Functional differences between AM and PR observed in the present study were surprisingly modest. In both patches, we found a large population of axis-tuned cells; the prominence of axis tuning in PR suggests that visual representation in perirhinal cortex is still partially feature-based rather than semantic. Furthermore, familiarity produced the same pattern of activity changes in both patches. Finally, in both patches we found a small population of super-sparse cells that responded most strongly to particular familiar individuals (**Fig. S12**, Supplementary Text), reminiscent of concept cells in the human medial temporal lobe (*39*). How can we reconcile these similarities with the differences that have been observed in previous studies (*33, 38, 40*)? Our experiments specifically probed visual familiarity and suggest that both (i) the tagging of stimuli as familiar, and (ii) the optimal separation of familiar identities, may largely be accomplished by IT cortex. However, it is likely that experiments probing visual association memory (including representation of degraded stimuli that require top-down feedback to complete) may reveal more prominent differences between AM and PR (*24, 30-33, 38*).

At present, due to the small number of faces we tested, we do not know if the distorted representations of familiar faces lie in an approximately linear subspace, in which case this distortion may be merely an affine transformation, or is situated on a more non-linear manifold, potentially consisting of distinct attractors for distinct familiar faces. Future experiments with a larger number of familiar faces will be needed to disambiguate these two scenarios.

Many brain areas across sensory, motor, and association cortex carry a representation of behaviorally relevant cognitive and sensory variables that is extremely low-dimensional compared to the number of neurons in the area (*3, 41, 42*). Such findings raise the question of why the brain is so large. The present study suggests that one possible reason may be to store memories of objects. By lifting representations of face memories into a separate subspace from that used to represent unfamiliar faces (**Fig. 3G**), attractor-like dynamics may be built around these memories to allow reconstruction of familiar face features from noisy cues (*43*) without interfering with veridical representation of sensory inputs. It is possible that different contexts invoke different memory subspaces (**Fig. S22**). To date, many studies of IT have emphasized the stability of response tuning over months (*44, 45*). Our results suggest such stability for representing unfamiliar stimuli co-exists with a precisely-orchestrated plasticity for representing familiar stimuli, through the mechanism of familiarity-gated change in axis.

## Author contributions

L.S. and D.Y.T. conceived the project and designed the experiments, L.S. and Y.S. collected the data, and L.S. and M.K.B. analyzed the data. L.S., M.K.B., and D.Y.T. interpreted the data and wrote the paper, with feedback from S.F. and Y.S.

## Data and materials availability

All data, code, and materials are available from the lead corresponding author upon reasonable request.

## Methods

Five male rhesus macaques (Macaca mulatta) of 5-13 years old were used in this study. All procedures conformed to local and US National Institutes of Health guidelines, including the US National Institutes of Health Guide for Care and Use of Laboratory Animals. All experiments were performed with the approval of the Caltech Institutional Animal Care and Use Committee.

### Visual stimuli

#### Face patch localizer

The fMRI localizer stimuli contained 5 types of blocks, consisting of images of faces, hands, technological objects, vegetables/fruits, and bodies. Face blocks were presented in alternation with non-face blocks. Each block lasted 24 s blocks (each image lasted 500 ms). In each run, the face block was repeated four times and each of the non-face blocks was shown once. A block of grid-scrambled noise patterns was presented between each stimulus block and at the beginning and end of each run. Each scan lasted 408 seconds. Additional details can be found in (*46*).

#### Monkey face model

To generate a large number of monkey faces, we built an active appearance model for monkey faces (*47*), similar to the method used for human faces in (*48*). Images of frontal views of 165 monkey faces were obtained from the following sources: a private database kindly provided by Dr. Katalin Gothard (101 images), the PrimFace database (visiome.neuroinf.jp/primface) (22 images), YouTube videos of macaques (19 images), and face images of macaques from our lab (23 images). The “shape” parameters were obtained by manually labelling 59 landmarks on each of the frontal face images (**Fig. S23A**). A 2D triangulated mesh was defined on these landmarks (**Fig. S23B**). The coordinates of the landmarks of each image were normalized by subtracting the mean and scaling to the same width, and a landmark template was obtained by averaging corresponding landmarks across faces. The “appearance” parameters were obtained by warping each face to the landmark template through affine transform of the mesh. To reduce the dimensionality of the model, principal component analysis was performed on both the coordinates of the landmarks (shape) and pixels of the warped images (appearance) independently. The first 20 PCs of shape and first 100 PCs of appearance were kept for the final model, capturing 96.1% variance in the shape distribution and 98.4% variance in the appearance distribution. We used this model not only to generate unfamiliar monkey faces, but also to compute shape-appearance features of familiar monkey faces (note: these faces were included in the 165-face database). For the latter, we projected the 59 landmarks and projected these onto the shape PCs; we then morphed the landmarks to the standard landmark template and projected the resulting pixels of the warped images onto the 100 appearance PCs.

#### Human face model

We built a human face model following the same procedure as for the monkey face model. Images of frontal views of 1200 human faces were obtained from different databases including: FERET (*49, 50*), CVL (*51*), MR2 (*52*), Chicago (*53*), and CelebA (*54*). Shape parameters used 92 landmarks (**Fig. S23C, D**).

#### Stimuli for electrophysiology experiments

Ten different sources of images were used to generate three different stimulus sets (**Fig. S2**).

1) Personally familiar human faces: Frontal views of faces of 9 people in the lab/animal facility who interacted with the subject monkeys on a daily basis.
2) Personally familiar monkey faces: Frontal views of faces of 9 monkeys in our animal facility that were current or previous roommates or cagemates of the subject monkeys, reconstructed using the monkey face model.
3) Personally familiar objects: Images of 8 toys the subject monkeys interacted with extensively.
4) Pictorially familiar human faces: Frontal views of faces of 8 people from the FEI database.
5) Pictorially familiar monkey faces: Frontal views of faces of 8 monkeys from the PrimFace database (visiome.neuroinf.jp/primface), reconstructed using the monkey face model.
6) Cinematically familiar human faces: Frontal views of faces of 18 main characters from 5 movies (Friends, The Big Bang Theory, The Mountain Between Us, Hard Candy, The Piano).
7) Cinematically familiar monkey faces: Frontal views of faces of 19 monkeys from 7 movies clipped from 7 videos from YouTube, reconstructed using the monkey face model.
8) Unfamiliar human faces: 1840 frontal view of faces from various face databases: FERET (*49, 50*), CVL (*51*), MR2 (*52*), Chicago (*53*), CelebA (*54*), FEI (fei.edu.br/∼cet/facedatabase.html), PICS (pics.stir.ac.uk), Caltech faces 1999, Essex (Face Recognition Data, University of Essex, UK; http://cswww.essex.ac.uk/mv/allfaces/faces95.html), and MUCT (www.milbo.org/muct). The background was removed, and all images were aligned, scaled, and cropped so that the two eyes were horizontally located at 45% height of the image and the width of the two eyes equaled 30% of the image width using an open-source face aligner (github.com/jrosebr1/imutils).
9) Unfamiliar monkey faces: 1840 images were generated using the monkey face model described above by randomly drawing from independent Gaussian distributions for shape and appearance parameters, following the same standard deviation as real monkey faces for each parameter. Faces with any parameter larger than 0.8 * maximum value found in a real monkey face were excluded to avoid unrealistic faces.
10) Unfamiliar objects: Images of objects were randomly picked from a subset of categories in the COCO dataset (arXiv:1405.0312). The choice of categories was based on two criteria: 1) only categories that our macaque subjects had no experience with (e.g., vehicles) were included, 2) categories with highly similar objects were excluded (e.g., stop signs). The included super-categories were: ‘accessory’, ‘appliance’, ‘electronic’, ‘food’, ‘furniture’, ‘indoor’, ‘outdoor’, ‘sports’, and ‘vehicle’. 1500 images of objects with area larger than 200^2^ pixels were isolated, centered, and scaled to the same width or height, whichever was larger.

We emphasize that due to the difficulty of obtaining a large set of high-quality monkey face images, we used the monkey face model described above to synthesize unfamiliar monkey faces; for consistency, all familiar monkey faces used in this study were also reconstructed using the monkey face model. Thus any differences in responses to familiar versus unfamiliar faces cannot be attributed to use of synthetic stimuli. Values of each feature dimension were normalized by the standard deviation of the feature dimension for analysis purposes.

From these 10 stimulus sources, four different stimulus sets were generated:

1) Screening set consisting of 8 or 9 images from 6 different categories (human faces, monkey faces, and objects, each either personally familiar or unfamiliar) (**Fig. S2A**). For unfamiliar stimuli, 8 novel images were used for each cell or simultaneously recorded group of cells. Each image was presented in random order, centered at the fixation spot, for 150 ms on 150 ms off (gray screen), repeated 5-10 times. The size of each image was 7.2° x 7.2°. Data using the screening stimulus set are shown in **Fig. 1, 4B, 4C, 4D1-3, 4E3, S3, S4A, S4B, S5, S6, S7, S18A, S19A, S20A-C**.
2) Thousand monkey face set consisting of 1000 unfamiliar (examples shown in **Fig. S2B**) and 36 familiar faces (personally familiar, pictorially familiar, and cinematically familiar faces, **Fig. S2A, C**), presented using the same parameters as the screening set, except for number of repetitions (3-5 times). In addition, the 8 novel unfamiliar faces shown in the screen set were shown again. Data using the stimulus set are shown in **Fig. 2, 3, 4D4, 4E1, 4E2, 4E4, S9, S10, S12, S15, S16, S17, S18, S20D-K, S21**.
3) Thousand human face set consisting of 1000 real human faces (examples shown in **Fig. S2B**), 35 familiar human faces, and 8 novel human faces (**Fig. S2A, C**). Other details same as for thousand monkey face set. Data using the stimulus set are shown in **Fig. S12**.
4) Face plane stimuli. Five stimulus sets each consisting of 10 x 10 human faces evenly spanning a 2D plane of face space ranging from −3 SD to 3 SD were generated by the human face model. The five face planes were randomly selected, and stimuli from the five planes were randomly interleaved during the experiment. Data using the stimulus set are shown in **Fig. S11**.

### Behavioral task

For electrophysiology and behavior experiments, monkeys were head fixed and passively viewed a screen in a dark room. Stimuli were presented on an LCD monitor (Acer GD235HZ). Screen size covered 26.0° x 43.9°. Gaze position was monitored using an infrared camera eye tracking system (ISCAN) sampled at 120 Hz.

#### Passive fixation task

All monkeys performed this task for both fMRI scanning and electrophysiological recording. Juice reward was delivered every 2-4 s in exchange for monkeys maintaining fixation on a small spot (0.2° diameter).

#### Preferential viewing task

Two monkeys were trained to perform this task. In each trial a pair of face images (7.2° x 7.2°) were presented on the screen side by side with 14.4° center distance (**Fig. 1B, S13**). Juice reward was given every 2-4 s in exchange for monkeys viewing either one of the images. Each pair of images lasted 10 s. Face pairs were presented in random order. To avoid side bias, each pair was presented twice with side swapped.

### MRI scanning and analysis

Subjects were scanned in a 3T TIM (Siemens, Munich, Germany) magnet equipped with AC88 gradient insert. 1) Anatomical scans were performed using a single loop coil at isotropic 0.5 mm resolution. 2) Functional scans were performed using a custom eight-channel coil (MGH) at isotropic 1 mm resolution, while subjects performed a passive fixation task. Contrast agent (Molday ION) was injected to improve signal/noise ratio. Further details about the scanning protocol can be found in (*55*).

#### MRI Data Analysis

Analysis of functional volumes was performed using the FreeSurfer Functional Analysis Stream (*56*). Volumes were corrected for motion and undistorted based on acquired field map. Runs in which the norm of the residuals of a quadratic fit of displacement during the run exceeded 5 mm and the maximum displacement exceeded 0.55 mm were discarded. The resulting data were analyzed using a standard general linear model. The face contrast was computed by the average of all face blocks compared to the average of all non-face blocks.

### Single-unit recording

Multiple different types of electrodes were used in this study. Single electrodes (Tungsten, 1 Mohm at 1 kHz, FHC) were used to collect most of the data. A Neuropixel prototype probe (128 channel, HHMI) was used to record ML from subject A. A multi-channel stereotrode (64 channel, Plexon S-probe) was used to record AM during muscimol silencing of PR in subject E. A chronic implanted microwire brush array (64 channel, MicroProbes) (McMahon, et al., 2014) was used to record from face patch AM in subject C. The electrode trajectories that could reach the desired targets were planned using custom software (*57*), and custom angled grids that guided the electrodes to the target were produced using a 3D printer (3D system). Extracelluar neural signals were amplified and recorded using Plexon. Spikes were sampled at 40 kHz. For single channel recorded data, spike sorting was performed manually by clustering of waveforms above a threshold in PCA space using a custom-made software (Kofiko) in Matlab. Multichannel recorded data was automatically sorted by Kilosort2 (github.com/MouseLand/Kilosort2) and manually refined in Phy (github.com/cortex-lab/phy).

### Muscimol experiment

To silence face patch PR, 1 µl (5 mg/ml) muscimol (Sigma) was injected into PR at 0.5 µl/min using G33 needle (Hamilton) connected to a 10 µl micro-syringe controlled by a micro-pump (WPI, UltraMicroPump 3). AM cells were recorded both before and 30 min after injection.

### Data analysis

All visually-responsive cells were included for analysis. To determine visual responsiveness, a two-sided T-test was performed comparing activity at [-50 0] ms to that at [50 300] ms after stimulus onset. Cells with p-value < 0.05 were included.

### Face selectivity index

A face selectivity index (FSI) was defined for each cell as:

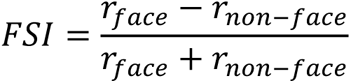

where *r* is the average neuronal response in a 50-300 ms window after stimulus onset (**Fig. S3**).

### Population average of response time course

For each cell, responses to the same stimulus category were first averaged in 10 ms time bins, then the responses were baseline-subtracted (using the average response in the time window 0 to 50 ms), and normalized by the maximum response across different stimulus categories after stimulus onset. The normalized responses were finally averaged across cells for each category after smoothing by a Gaussian function with 10 ms standard deviation (**Fig. 1E** right, **Fig. 3B, S4B, E, S20C, I**).

To determine the time point at which responses rose above baseline (e.g., **Fig. 1E**), we compared the response at each time point to the baseline response (average response over [-50 0] ms) using a one-tailed T-test, and determined the first time point at which P < 0.01.

### Preferred axis of cells

The preferred axis of cells was computed in two different ways:

#### Spike-triggered average (STA)

The average firing rate of a neuron was computed to each stimulus, either in a full time window [50-300] ms or sliding 50 ms time window after stimulus onset. The STA was defined as:

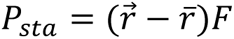

where 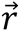 is *1 × n* vector of the firing rate response to a set of *n* face stimuli, 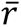 is the mean firing rate, and *F* is a *n × d* matrix, where each row consists of the *d* parameters representing each face stimulus in the feature space.

#### Linear regression/Whitened STA

For a small sample of stimuli, e.g., 36 familiar faces, the features are not necessarily white (i.e., uncorrelated). As a control, to ensure that the difference in STA observed in **Fig. 2B, D** was not due to mismatched feature distributions between familiar and unfamiliar faces, we repeated our main analysis using a whitened STA (**Fig. S15B**) as follows:

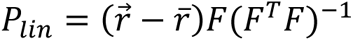

For all figures except **Fig. S11**, we used 20 dimensions to compute the preferred axis (first 10 shape and first 10 appearance dimensions). For **Fig. S11**, we used two dimensions (the two dimensions spanning each of the five face planes).

### Principal orthogonal axis

The principal orthogonal axis was defined as the *longest* axis orthogonal to its preferred axis. First, for each of the 1000 unfamiliar face images represented as *d*-dimensional vector 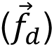 in face feature space, its component along the preferred axis (*P*) of the cell was subtracted

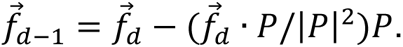

Then principal component analysis was performed on the set of 1000 vectors 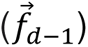, and the principal orthogonal axis was the first principal component.

### Quantifying significance of axis tuning

For each cell, we compared the explained variance by the axis model to a distribution of explained variances computed for data in which stimulus identities were shuffled (1000 repeats). We considered axis tuning significant if the frequency of a higher explained variance in the shuffle distribution was less than 5% (**Fig. S9A, C, E**).

### Quantifying consistency of preferred axis

For each cell, the stimuli were randomly split into two halves, and a preferred axis was calculated using responses to each subset. Then, the Pearson correlation (*r*) was calculated between the two. This process was repeated 100 times, and the consistency of preferred axis for the cell was defined as the average *r* value across 100 iterations (**Fig. S9B, D, F**).

### Face feature decoding and reconstruction

To decode face features, firing rates after stimulus onset in a chosen time window (see **Fig. 2E** legend) were first averaged across multiple repeats of the same stimulus, then linear regression was performed on a training set of 999 unfamiliar faces to compute the linear mapping from population response vector 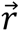 to face feature vector 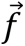:

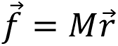

The decoding was performed on the remaining one unfamiliar and all familiar faces using this mapping *M*. Decoding accuracy was measured by (i) the correlation coefficient between decoded and actual face features (**Fig. 2E1**), (ii) the mean square error between decoded and actual face features (**Fig. 4E4**), or (iii) the rate of correctly choosing the actual face shown as the decoded face among a given number of randomly-sampled distractor faces (**Fig. S9H**); for the last method, the decoded face was selected as the face with minimum Euclidean distance in feature space to the decoded feature vector. For all three methods, the decoding accuracy for unfamiliar faces was computed 1000 times through leave-one-out cross validation.

To reconstruct faces (**Fig. 2E2**), we built a face feature decoder using responses to unfamiliar faces, computed either in a short ([120 170] ms) or long ([220 270] ms) latency window.

### Face identity decoding

To decode face identity (**Fig. 2G**), firing rates after stimulus onset in a chosen time window of each trial were randomly split in half and averaged. Then a multi-class linear SVM decoder was trained to classify each face identity for 30 familiar or 30 unfamiliar feature-matched (**Fig. S14**) faces separately, using one half for training and testing on the other half. This was repeated 20 times.

### Testing the contribution of axis change to face distance increase

We compared mean pairwise distances between the 30 familiar faces (**Fig. S14**) by predicting, in a cross-validated way, responses to each familiar face pair using two different axes for each cell (**Fig. 2H2**): (i) the familiar axis, (ii) the unfamiliar axis. Specifically, for each cell, the axis direction was computed using long latency (150-300 ms) responses to the 28 remaining familiar or unfamiliar feature-matched faces, and normalized to have unit length; for both familiar and unfamiliar axes, the gain was fit using responses to the 28 remaining familiar faces. This yielded four vectors, ****pred***_familiar/unfamiliar a*x*is_f*a*ce_1/2_*. For each face pair, we then computed the Euclidean distances *d(***pred***_familiar a*x*is_f*a*ce_1_, ***pred***_familiar a*x*is_f*a*ce_2_)*, and d(****pred***unfamiliar a*x*isf*a*ce*_1_, ****pred***unfamiliar a*x*isf*a*ce*_2_).

### Sparseness

The sparseness of a cell (**Fig. S12C**) was measured by Gini coefficient defined by:

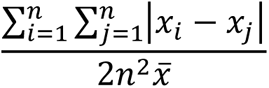

where *x_i_* is the firing rate response to *i*th stimulus, *n* is total number of stimuli, and 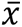 is mean firing rate to all stimuli. If responses to all stimuli are equal, the Gini coefficient is 0. If the response to one stimulus is 1 and responses to all other stimuli are 0, the Gini coefficient is 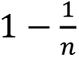.

### Response reliability

The response reliability of a cell (**Fig. S16B**) was computed as the Pearson correlation coefficient between the average firing rate of half the trials (50-300 ms after stimulus onset) and that of the other half (randomly split), using responses of the cell to the thousand face set.

### Familiarity decoding

Firing rates after stimulus onset in a chosen time window (stated for each particular case in the figure legends) were first averaged across multiple repeats of the same stimulus, then the decoding accuracy was obtained as the average of leave-one-out cross-validated linear SVM decoding. For the thousand face set, the training sample was balanced by randomly subsampling 36 unfamiliar faces, repeated 10 times (**Fig. 3C**).

### Centroid shift analysis

To determine the time when the shift of the neural representation centroids for familiar and unfamiliar faces provided familiarity discriminability (**Fig. 3C**), population responses (50 ms sliding time window, step size 10 ms) to all 36 familiar faces and randomly subsampled 36 unfamiliar faces were first projected to the axis connecting neural centroids of familiar and unfamiliar faces. Then *d’* was computed for the projected values:

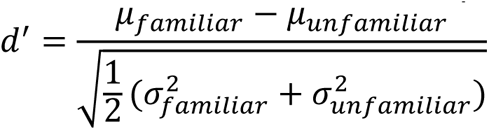

Here *µ* and *σ* are the mean and variance of the projected values, respectively. The computation was repeated 10 times for each random subsampling of 36 unfamiliar faces. Chance level d’ was estimated by randomly shuffling the population responses to each face 10 times. The time when d’ was significantly higher than chance was determined by one-tailed T-test (p<0.01).

### Analysis of orthogonality between familiarity and face feature decoding axes

To determine the cosine similarity between familiarity and face feature decoding axes (**Fig. 3F**), we obtained the familiarity decoding axis as described above in the section on “Familiarity decoding.” For unfamiliar faces, we obtained the face feature decoding axis as described in the section above on “Face feature decoding and reconstruction.” For familiar faces, we obtained the face feature decoding axis by computing the pseudo-inverse of the face feature encoding axis (necessary due to the small number of familiar faces); we obtained the latter as described above in the section on “Preferred axis of cells” (using the STA).

### Computing normalized firing rate changes

Normalized firing rate change in **Fig. 4D1** was computed as follow:

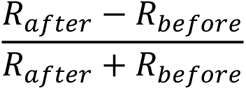

where *R_before_* is the mean firing rate within 50-300 ms after stimulus onset before Muscimol injection, and *R_after_* is the same for after muscimol injection.

### Matching face feature distributions

We wanted to ensure that the difference in preferred axis (**Fig. 2B, 2D**) and the difference in pairwise distance in the neural state space (**Fig. 2F-H**) were not due to mismatched feature distributions between familiar and unfamiliar faces. To this end, we identified a *feature-matched subset* of 30 familiar and 30 unfamiliar faces. For the top 20 face features, these two face sets were matched in feature variance **(Fig. S14A)**, distribution of pairwise face distances in feature space **(Fig. S14B)**, and distribution of each feature **(Fig. S14C)**. This was achieved by searching for a subset of faces that minimized the following cost function:

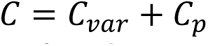

The first term evaluated the difference of variance:

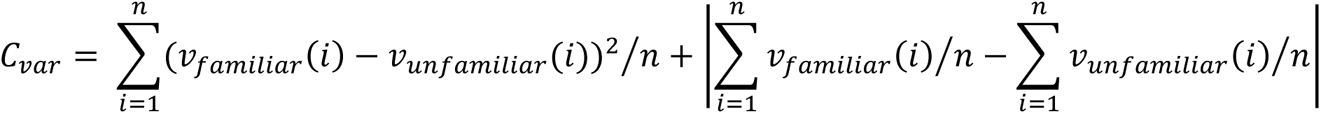

where *v(i)* is the variance of the *ith* feature, *n* = 20 is the number of features. It is the sum of mean square error and absolute value of mean difference between the variance of each feature.

The second term ensured the distributions in consideration are not significantly different, which was measured by the *p* values of K-S test being larger than 0.05:

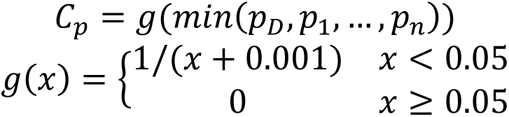

where *p*_*D*_ is the *p* value of K-S test between distributions of pairwise face distances for familiar vs. unfamiliar, *p*_*i*_ is the *p* value of K-S test between distributions of the *ith* feature for familiar vs. unfamiliar.

The optimization was performed using a gradient-descent-like algorithm: in each iteration *dC* was estimated by removing or adding each face, and the change that decreased *C* the most was applied, until *C* did not decrease anymore. To balance the number of familiar and unfamiliar faces in the result, we set a minimum number of familiar faces (23–36). When the number was chosen to be 30, the resulting number of unfamiliar faces also happened to be 30.

Finally, we confirmed that for the resulting set of 30 familiar and 30 unfamiliar faces, the faces were indeed feature matched (**Fig. S14**), and the axis model explained similar amounts of variance for both familiar and unfamiliar face responses (**Fig. S9E, F**).

In **Fig. S2B**, to demonstrate the diversity of faces in the 1000 face set and the capability to match each of our familiar face sets, we used the same matching method with different subsets of familiar faces.

## Supplementary Text

### Is the feature tuning of face cells ramp or V-shaped?

A recent study (*58*) reports that cells in face patch AM show prominent V-shaped tuning around the average face at long latencies (peaking at ∼200 ms), and suggests that this V-shaped tuning may have been missed by earlier studies (*48, 59*) because they did not use a long enough presentation time (300 ms). In the main experiments of the present study, all visual stimuli for electrophysiology experiments were presented for 150 ms ON and 150 ms OFF.

To determine the extent to which presentation time affects our results, for a subset of cells, we repeated our experiment to map the preferred axis of cells (**Fig. 2**) using a stimulus presentation time of 300 ms ON and 300 ms OFF, exactly matching the presentation time used by Koyano et al. (*58*). We varied faces along two axes within 5 randomly chosen planes, such that a 10×10 grid was sampled within each plane (**Fig. S11A**). This ensured that the average face was presented as part of our stimulus set, to more closely mimic the stimulus used by Koyano et al. (in our main experiment, we sampled faces from a 120d face space by sampling each dimension from an independent Gaussian distribution; with this approach, the average face has very low probability of being presented, as almost all of the stimulus density is on an outer shell of the feature space).

We continued to observe exclusively ramp-shaped tuning (**Fig. S11B, C**). We speculate that the V-shaped tuning observed by Koyano et al. may have been due to greater familiarity of stimuli closer to the average face due to the stimulus design used in that study: 12 morph trajectories were presented, each crossing the same average face, resulting in 145 total stimuli; across the 12 trajectories, stimuli near the average all looked very similar (∼4*12/145 = 33% of stimuli). In contrast, because (i) we sampled over a 2D face space, and (ii) our extreme faces were more caricatured (3 S.D. from the average), the proportion of faces similar to the average was much smaller in our stimulus set, ∼2*2/100 = 4% (**Fig. S11A**). This interpretation of Koyano et al.’s finding is consistent with the fact that the reduced responses to stimuli at the trough of the V observed by Koyano et al. occurred at increased latency, matching the time course of suppression for familiar stimuli observed in our first experiment (**Fig. 1E**). As noted in the main text, this suppression to familiar faces depends strongly on temporal context (**Fig. 3A, S18A**). Importantly, it occurs for multiple distinct familiar faces (**Fig. 1E**), and hence is unlikely to be solely a form of predictive normalization of incoming stimuli.

### A small subset of AM and PR cells respond extremely sparsely to familiar faces

In addition to cells with ramp-shaped tuning, we found a small population of cells that responded extremely sparsely to just a few of the stimuli (**Fig. S12**). PR contained significantly more such super-sparse cells than AM (**Fig. S12**; sparsity > 0.7, p < 0.015, Chi-square test, degree of freedom = 1). The frequency of preferred stimuli being familiar faces was much higher than chance: in PR (AM), for 8/11 (2/5) cells with sparsity > 0.7, the most effective stimulus out of 1000 unfamiliar faces and 36 (monkey) or 35 (human) familiar faces was a familiar face. The probability that this would happen by chance is ∼(11 choose 8)(36/1036)^8^(1000/1036)^3^ = 3.2*10^-10^. Thus these super sparse cells are clearly specialized for representing familiar faces. Their extraordinary specificity of response to specific familiar individuals is reminiscent of “concept cells” found in the human temporal lobe. Future work may test whether these cells respond not just to the face of a familiar individual but also to sounds and symbols evoking that individual, suggesting a representation of abstract concept (*60*).

The fact that PR contains both axis- and exemplar-tuned cells suggests that it may inherit the AM axis representing physical identity and then compute a transformation of this code to an exemplar-based code more suitable for representing mnemonic associations between specific familiar stimuli (*61–63*).

### A hierarchical generative inference model predicts gain change from PR inactivation

What could be the function of the gain changes induced by PR? A prominent model of vision asserts that the visual system implements a hierarchical inference network in which stimuli are discriminated in a feedforward pathway and predictively generated in a feedback pathway (*64–67*). In such a network, when the input stimulus is ambiguous, top-down feedback serves to fill in missing information based on prior knowledge. When the input stimulus is unambiguous (as was the case for all of our stimuli, which consisted of clear, high resolution faces), then the “feedback receptive field,” namely, the selectivity of a cell to stimuli resulting from neural activity in the feedback generative pathway, should be identical to the feedforward receptive field (**Fig. S19B-D**). Thus in a hierarchical inference network, inactivating feedback should simply change the gain of responses to clearly visible stimuli, matching our experimental results. Further experiments, e.g., with ambiguous stimuli, would be needed to prove that PR feedback subserves hierarchical inference.

### A simple model of face patch representations

Here we will briefly describe a straightforward network model that recapitulates some of the basic features of the neural representations we have described in our paper. We formulate this model as a population of firing rate neurons with sigmoidal non-linearities which receive inputs that encode the features of the faces presented to the system as stimuli. Specifically, we model the input to each face patch neuron as a linear combination of the face features with coefficients that are chosen randomly and independently from a Gaussian distribution, in addition to bias (offset) and noise terms. If the activations of the model face patch neurons in response to the presentation of unfamiliar faces remain largely restricted to the linear regime of the sigmoid, then since the input currents are confined to a linear subspace (of dimensionality equal to number of encoded face parameters) the neural activities will also be located on an approximately linear subspace of the firing rate space (modulo noise). In this way we can achieve an approximately linear encoding of the face features, by embedding the manifold of face parameters in an almost linear fashion in the firing rate space of the neural population.

We would like to qualitatively reproduce the basic coding properties of the recorded neural data using this model. At short latency (after stimulus presentation), the main observed difference between unfamiliar and familiar face representations is a relative shift in the centroids of their respective coding manifolds. We can easily introduce such a shift by adding to the input current of each neuron a bias term (again drawn randomly from a Gaussian distribution) whose sign depends on the familiarity of the stimulus. Repeating the analysis of the cosine similarity between the preferred axes of cells in response to unfamiliar and familiar faces as in **Fig. 2B**, we find that its distribution is essentially identical to the control (**Fig. S21B**) (which is the cosine similarity between unfamiliar preferred axes and those computed on a small subset of unfamiliar faces). This is consistent with the time-resolved analysis of **Fig. 2D**, which shows no significant differences at short latency. The shift between familiar and unfamiliar coding manifolds allows a linear decoder to discriminate familiar from unfamiliar faces in a way that generalizes to held-out stimuli (i.e., faces not used during training), again in agreement with **Fig. 3**.

For the long latency response to face stimuli, we observed in the data that the preferred axes corresponding to familiar and unfamiliar stimuli are less correlated (while those in response to unfamiliar stimuli are quite consistent, i.e., show a large cosine similarity with the axes computed on a subset of unfamiliar faces) (**Fig. 2B**). To capture this effect, we consider that the input to the recorded population of neurons may arrive through two different pathways (which can both still be modeled with random, independently chosen Gaussian weights) that are modulated by the familiarity signal in different ways (see schematic in **Fig. 21A**). This familiarity signal was already decodable from the neural activity at short latency due to the shift discussed above (which is still present at long latency). However, while we considered both input pathways to be activated to the same extent by familiar and unfamiliar faces at early times, we now let unfamiliar faces activate only one of these pathways, and familiar faces exclusively but strongly activate the other pathway at longer latency. This procedure reproduces the small correlation between the axes for familiar and unfamiliar faces seen in **Fig. 2B**.

The strong input current for familiar faces leads to responses that have a larger variance (**Fig. 21C**). This larger variance leads to an increased pairwise distance between familiar faces as compared to the distance between pairs of unfamiliar faces, consistent with **Fig. 2F**. While we do not explicitly model axis change here (as observed in the actual data, **Fig. 2B, D, E, H**), it would provide another mechanism to increase distances between pairs of familiar faces, if axis change leads to better alignment with directions of maximum feature variance in the familiar faces. Familiarity can still be decoded, as in the earlier time interval, but in addition these long latency representations also allow us to successfully discriminate particular familiar faces from others with high accuracy.

---

**Figure S1.**
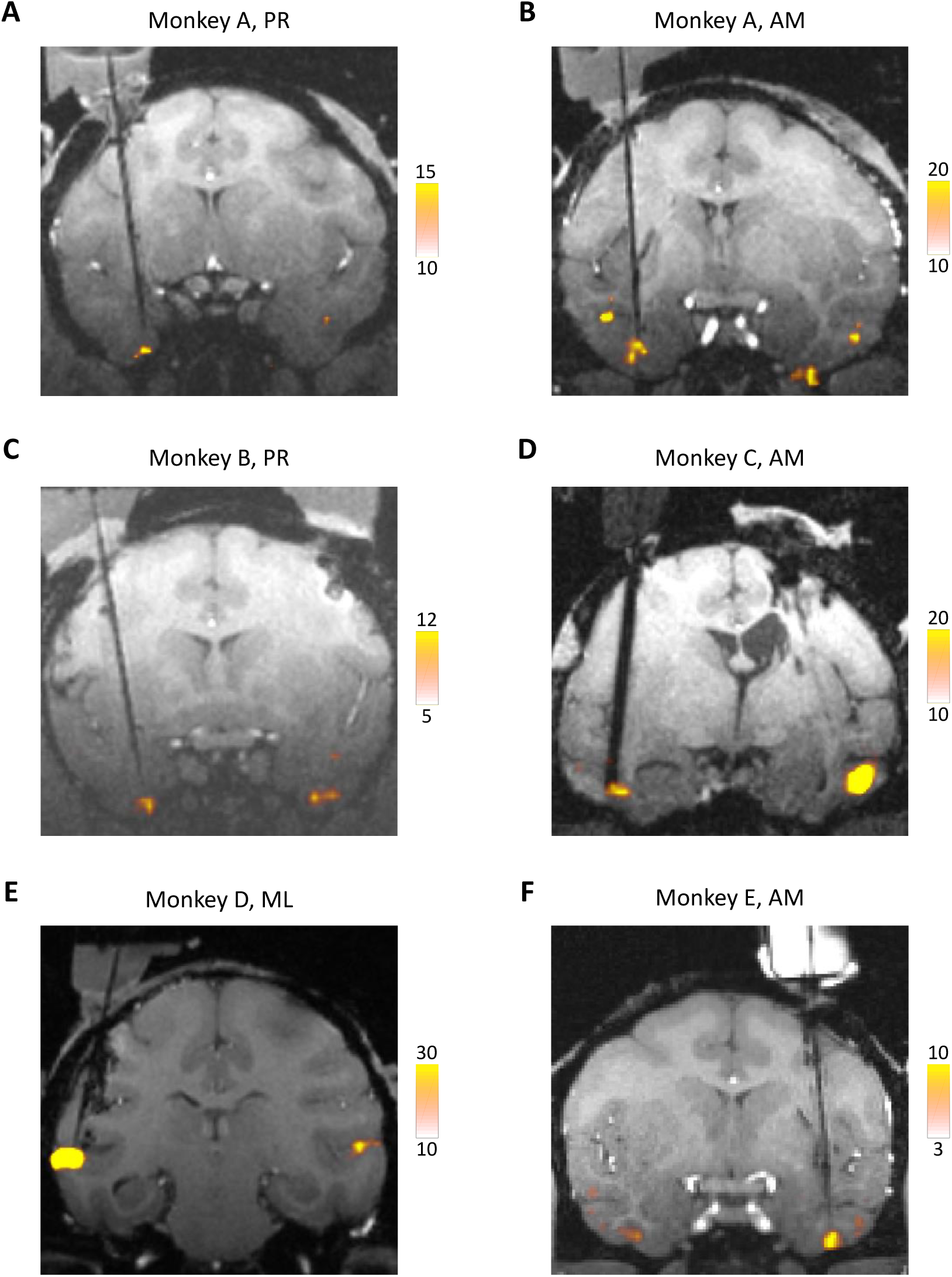
Coronal slices showing the electrode targeting six recording sites from 5 monkeys. **(A)** Single electrode targeting PR in monkey A. **(B)** Single electrode targeting AM in monkey A. **(C)** Single electrode targeting PR in monkey B. **(D)** Brush array electrodes targeting AM in monkey C. **(E)** Single electrode targeting ML in monkey D. **(F)** Single electrode targeting AM in monkey E. Activations for the contrast faces versus objects are shown, at uncorrected p values.

**Figure S2.**
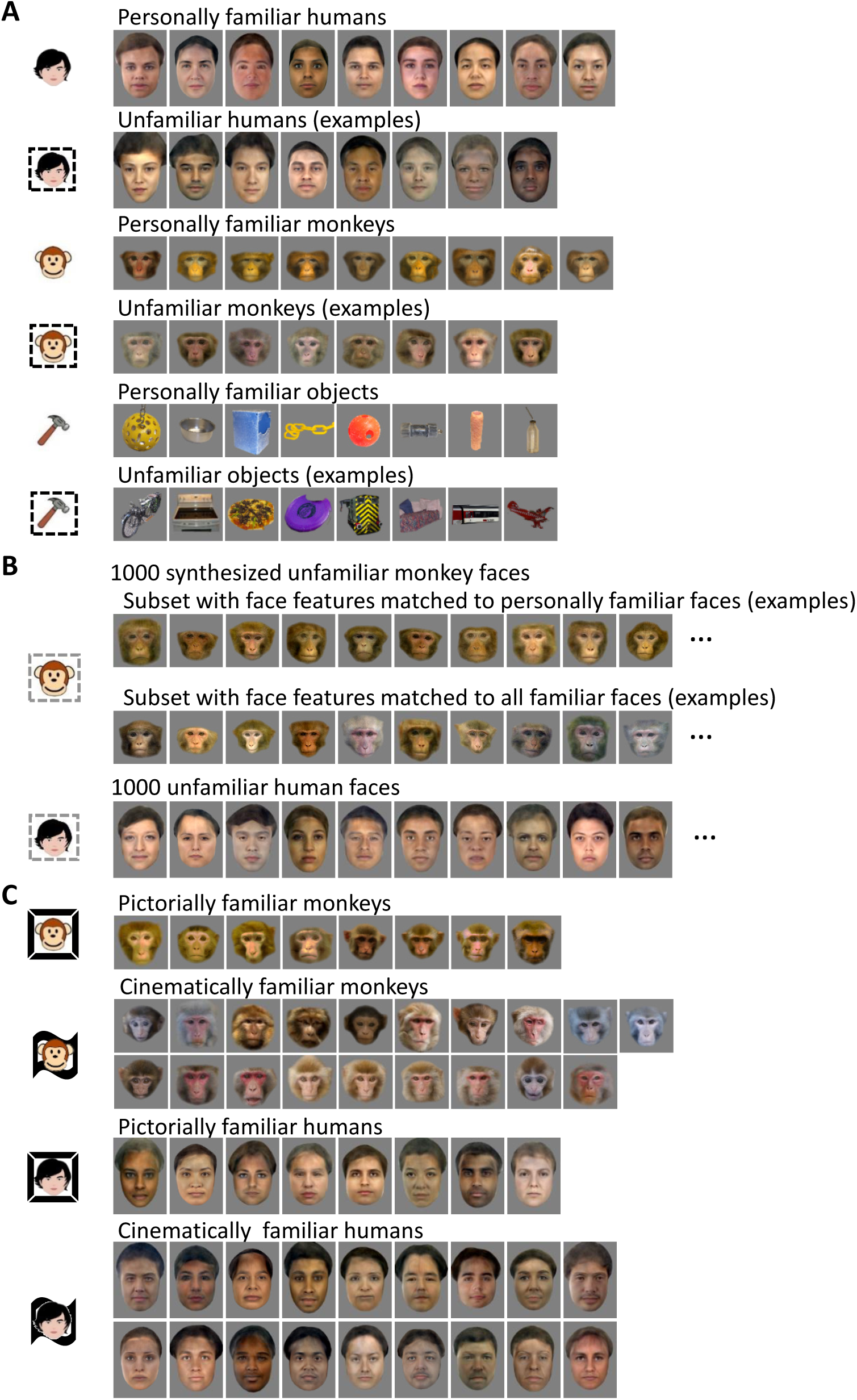
Visual stimuli. **(A)** Screening stimuli. Example unfamiliar stimuli are shown here; a new set was presented for every recording site, drawn from image sets described in the Methods. **(B)** Examples of unfamiliar faces in the thousand face stimulus set. Monkey faces were generated by a 120d shape-appearance model (see Methods). The thousand monkey face stimulus set was extremely diverse, allowing subsets of faces to be chosen that were matched in feature distributions to familiar faces (see Methods). Shown here are examples from two subsets, one matched to the personally familiar faces, and one matched to all familiar faces. Human faces (used for analysis in Supplementary Text) were from online databases. **(C)** Additional familiar faces (pictorially and cinematically familiar). [Note: all human faces in this figure have been replaced by synthetically generated faces due to biorxiv policy on displaying human faces.]

**Figure S3.**
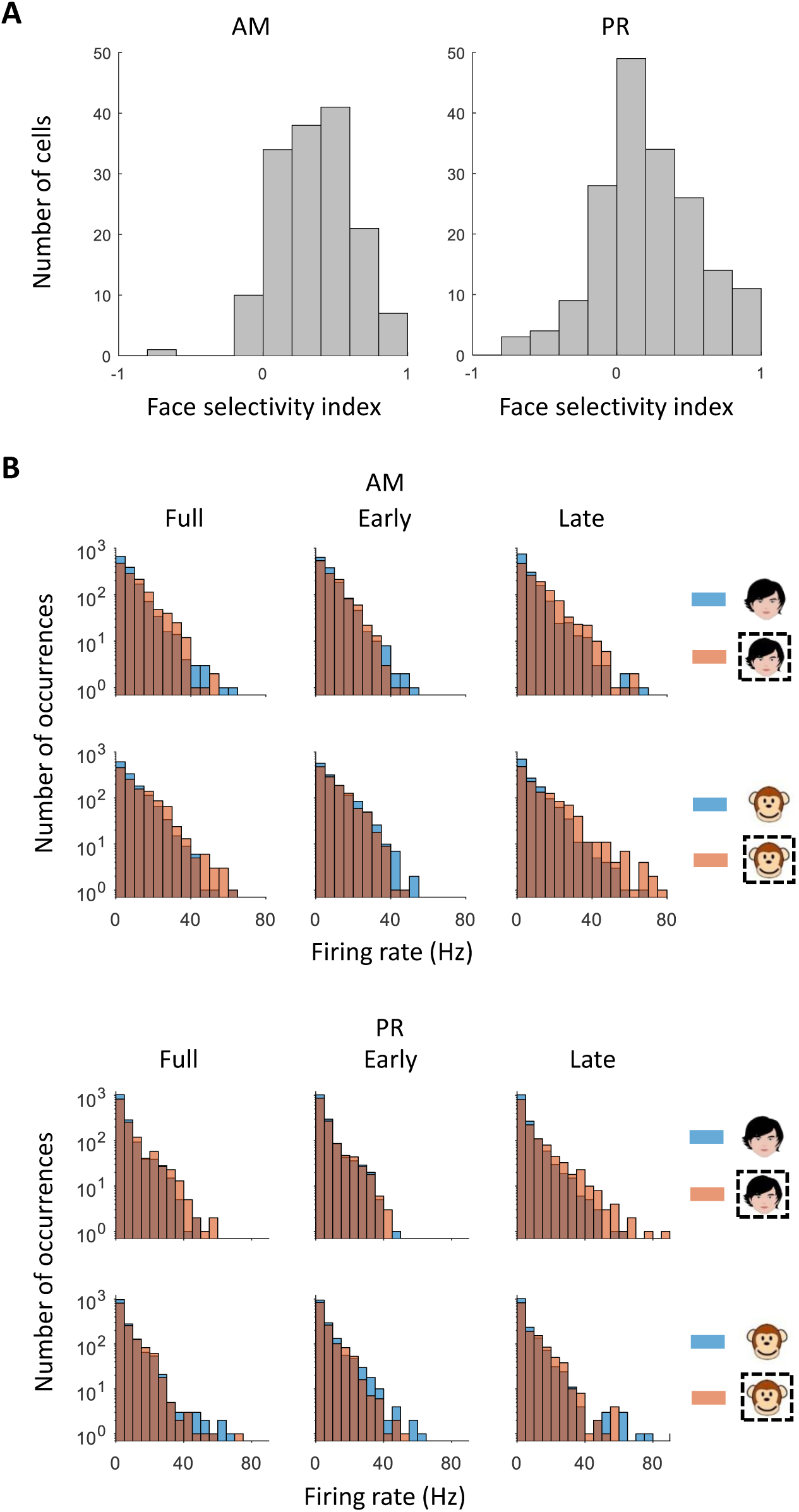
**(A)** Histograms of face selectivity indices computed using screening stimuli from AM and PR (see Methods). **(B)** Response distributions from AM and PR to screening stimuli for familiar or unfamiliar faces at three different time windows. Icon conventions as in Fig. 1C.

**Figure S4.**
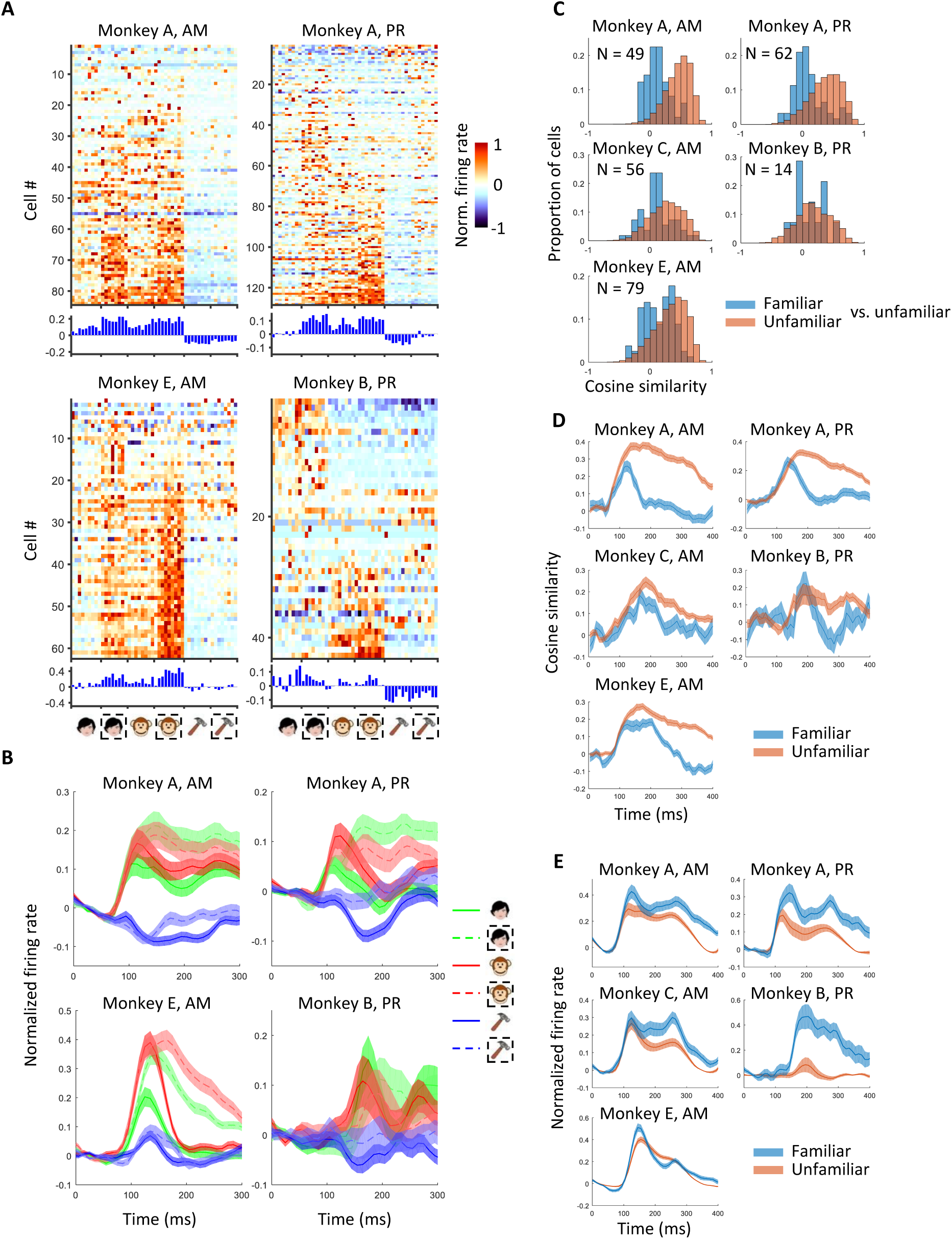
Main results computed separately for each animal individually. **(A)** Responses of cells to stimuli from six stimulus categories (same as Fig. 1**C1**). **(B)** Response time course averaged across cells and exemplars within each screening category (same as Fig. 1E, right). **(C)** Population analysis comparing preferred axes for familiar versus unfamiliar faces (same as Fig. 2B). **(D)** Time course of the similarity between preferred axes for unfamiliar-unfamiliar (orange) and unfamiliar-familiar (blue) faces (same as Fig. 2D). **(E)** Average response time course for 1000 monkey face stimulus set (same as Fig. 3B).

**Figure S5.**
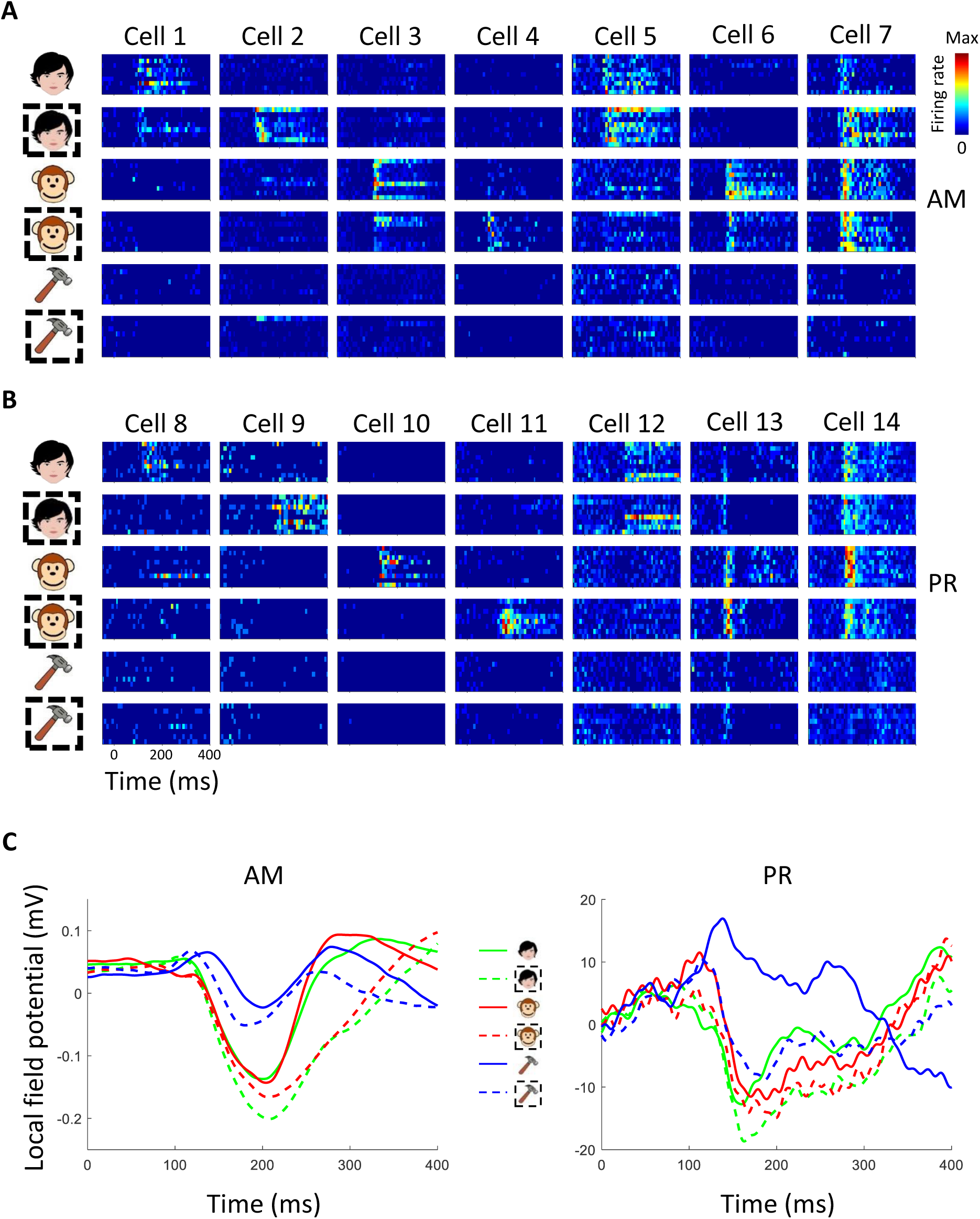
Responses of example neurons to familiar and unfamiliar screening stimuli. Seven example cells from AM. Icon conventions as in Fig. 1. **(B)** Seven example cells from PR. **(C)** Local field potentials from an example site in AM and one in PR.

**Figure S6.**
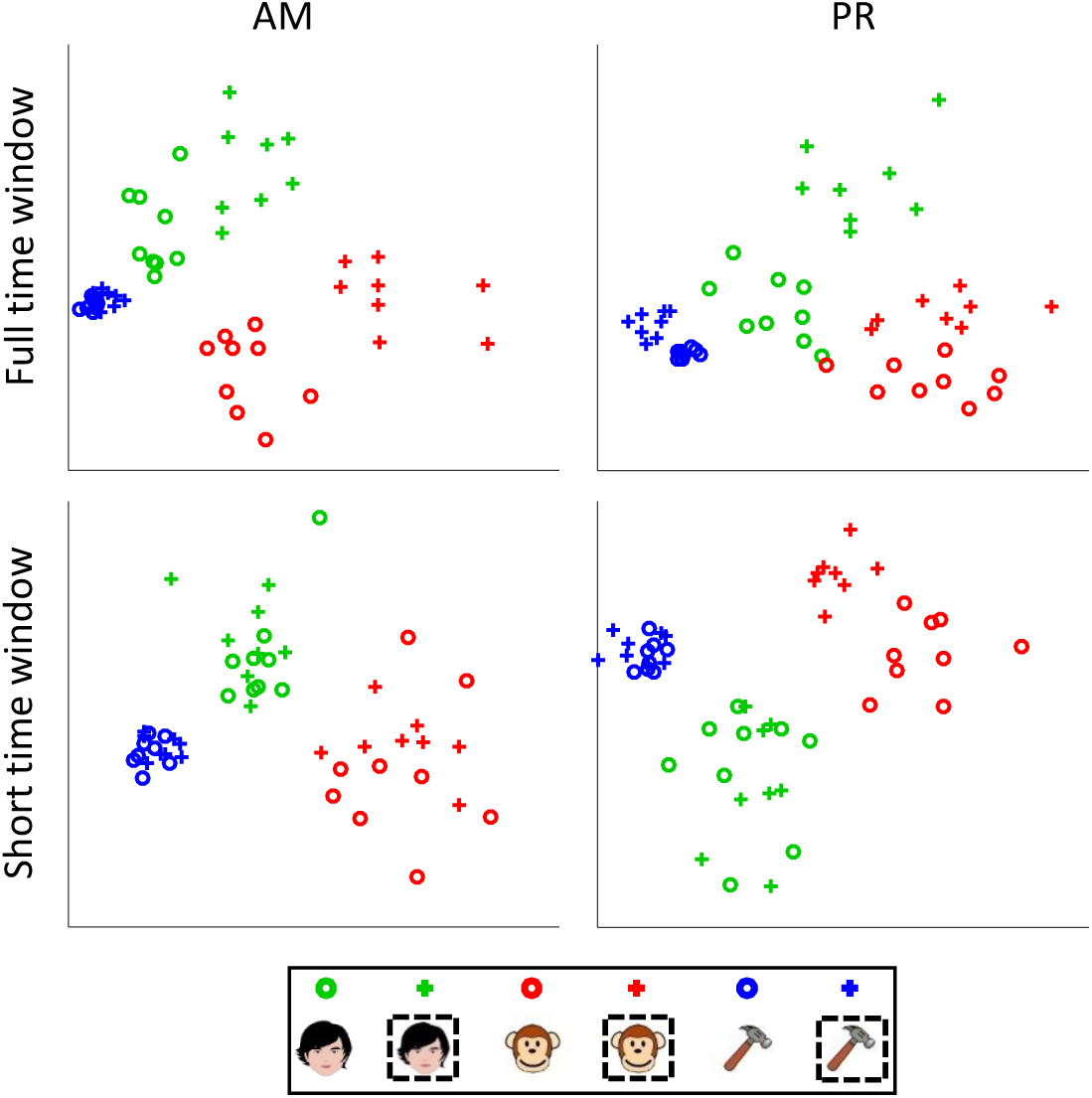
MDS analysis for AM and PR. Top: full time window, bottom: short time window. Time windows as in Fig. 1C. The percentage of total variance captured by the two dimensions was 50.7% (AM, full), 46.4% (AM, short), 35.5% (PR full), 41.9% (PR short).

**Figure S7.**
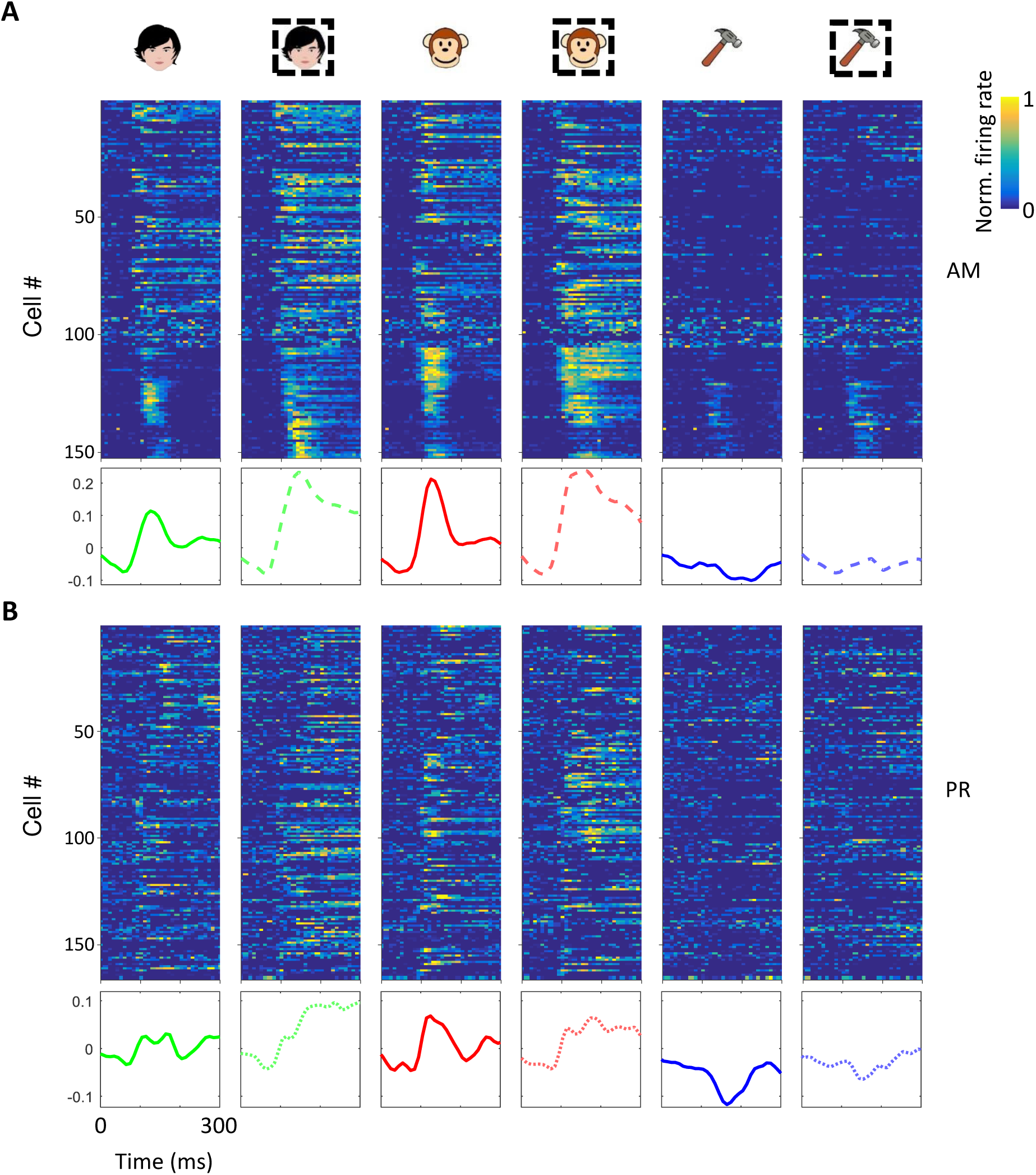
Response time courses from AM and PR. **(A)** Top: response time courses of individual neurons from face patch AM. Bottom: response time course averaged across neurons. Icon conventions as in Fig. 1C. **(B)** Same as (A) for PR.

**Figure S8.**
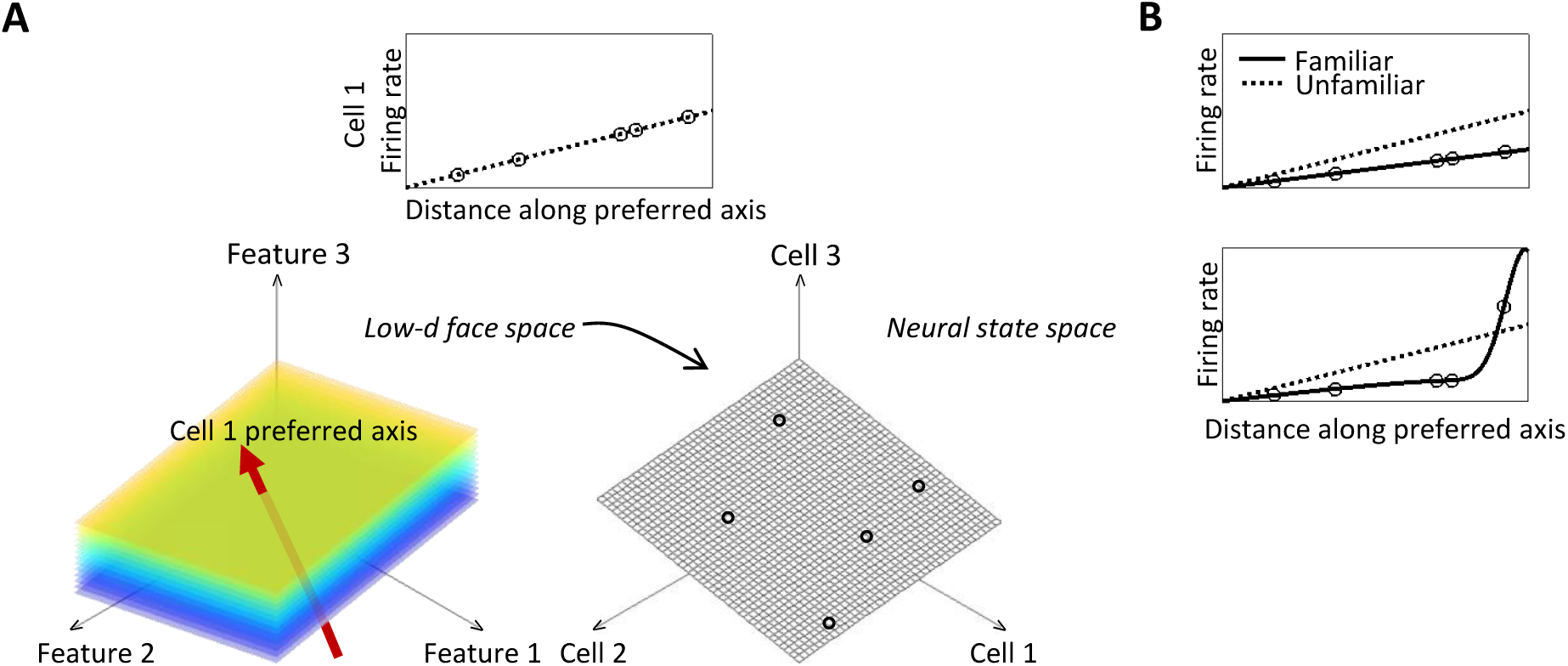
Geometry of axis coding. **(A)** Schematic of axis code used by face patches to represent facial identity. Top: Response of an example axis-tuned cell along its preferred axis. Left bottom: Response of the cell as a function of location in the face feature space. Colors indicate response magnitude; red arrow indicates preferred axis. Right bottom: the face feature space is embedded in the neural state space as a linear subspace. Circles indicate responses to a specific subset of faces. The axis code implies that responses to unfamiliar faces are confined to this subspace. **(B)** Familiarity has previously been proposed to produce a monotonic transform of responses such as scaling (top) or sparsening (bottom).

**Figure S9.**
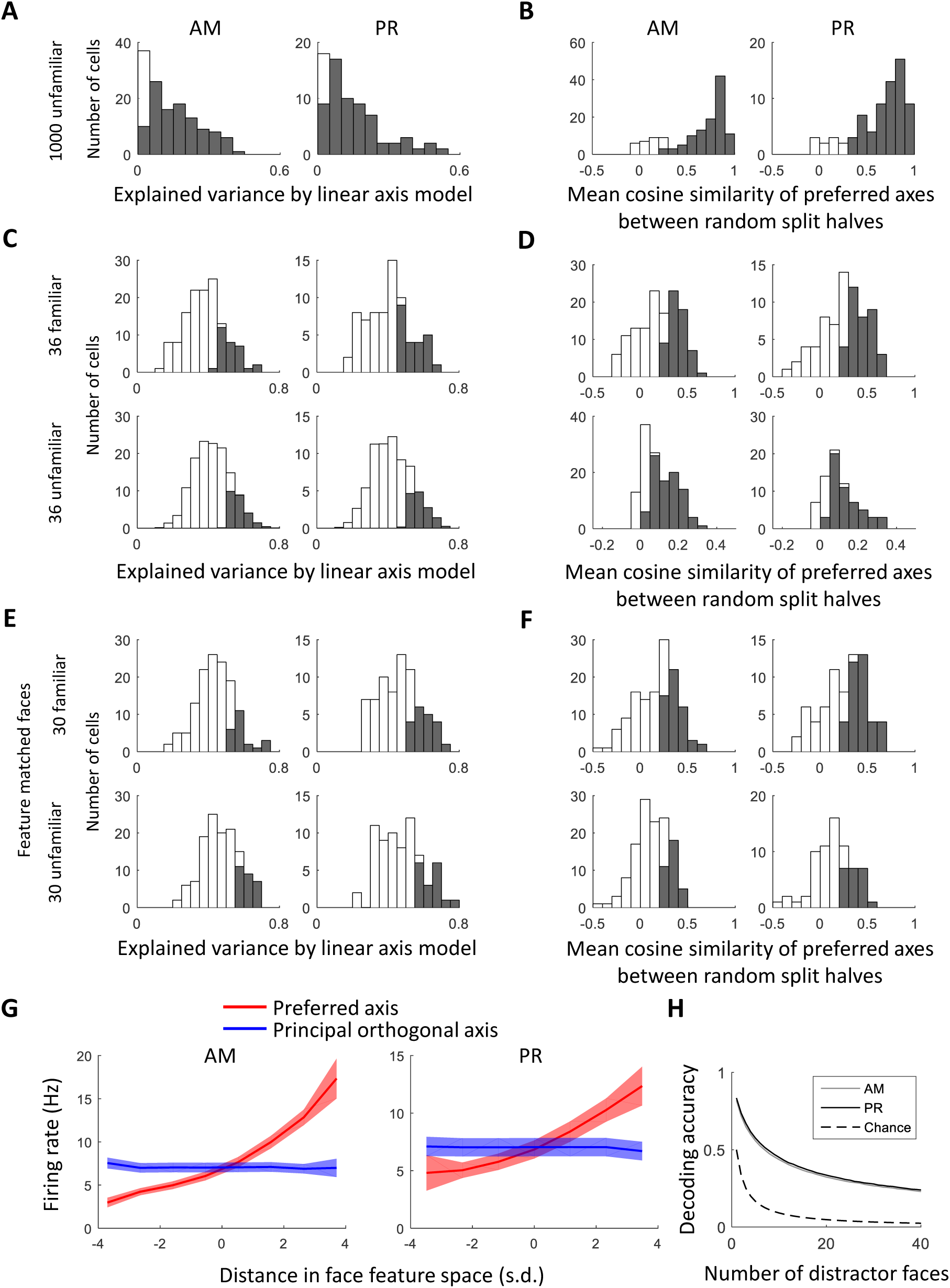
Quantification of axis tuning in AM and PR. **(A)** Distribution of explained variance by the linear axis model for responses to 1000 unfamiliar faces; shaded bars indicate the subset of cells for which the explained variance was significantly higher than for stimulus-shuffled data (1000 repeats). **(B)** Distributions of mean cosine similarity of preferred axes across repeated split halves (100 repeats) of responses to 1000 unfamiliar faces for AM and PR. Same conventions as in (A). **(C, D)** Same as (A) and (B) but for 36 familiar faces or 36 randomly-sampled unfamiliar faces (20 repeats). **(E, F)** Same as (A) and (B) but for 30 familiar or 30 unfamiliar feature-matched faces. **(G)** Red lines show the average modulation along the preferred axis across the population of AM and PR cells. Blue lines show the average modulation along the longest axis orthogonal to the preferred axis in the 50d face space that accounts for the most variability. Shaded area, SEM. **(H)** Comparison of face decoding performance for AM versus PR using a leave-one-out cross validated linear decoder as a function of number of random distractor faces (see Methods).

**Figure S10.**
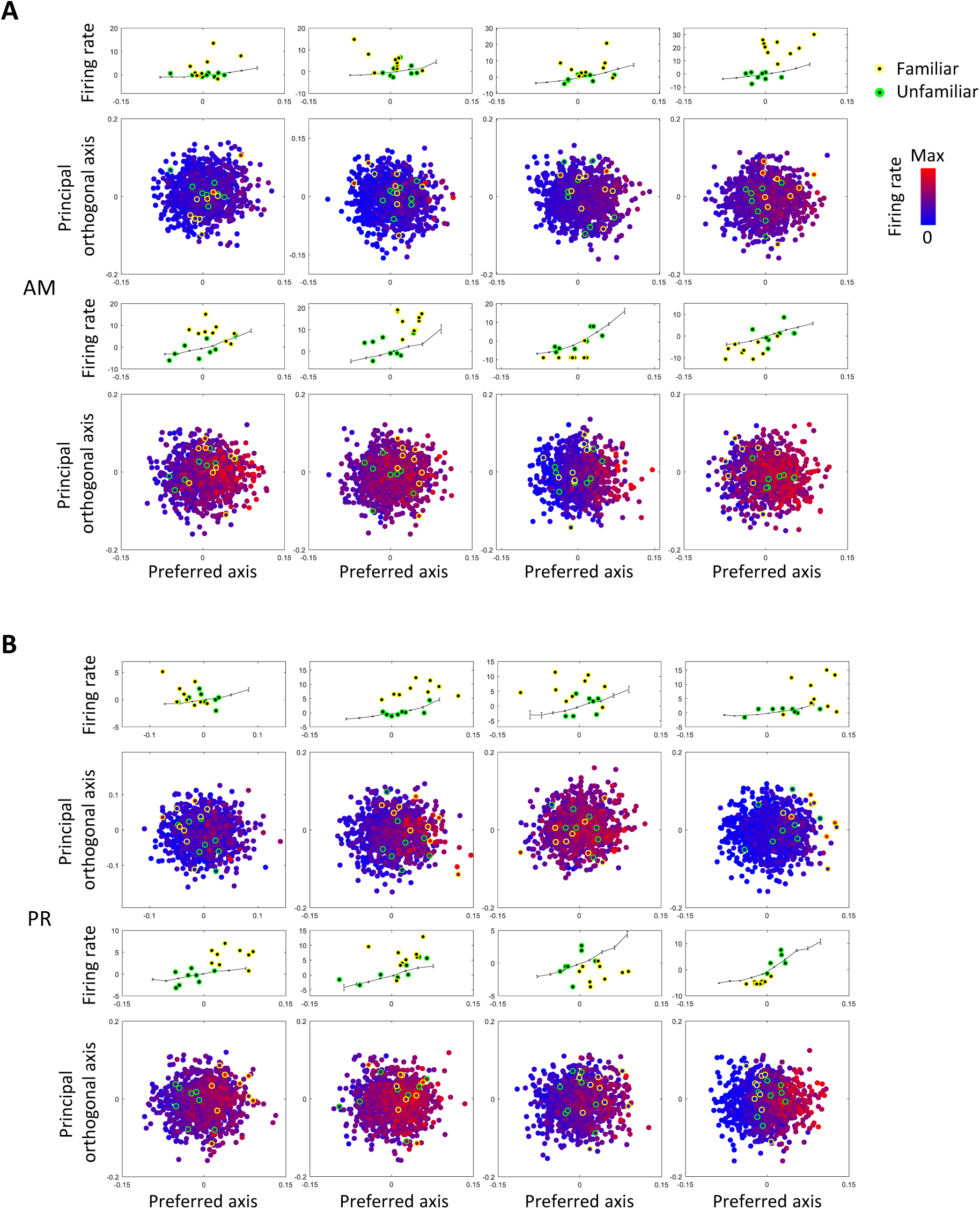
Examples of additional axis-tuned cells. **(A)** 8 cells from AM (same format as Fig. 2A). **(B)** 8 cells from PR.

**Figure S11.**
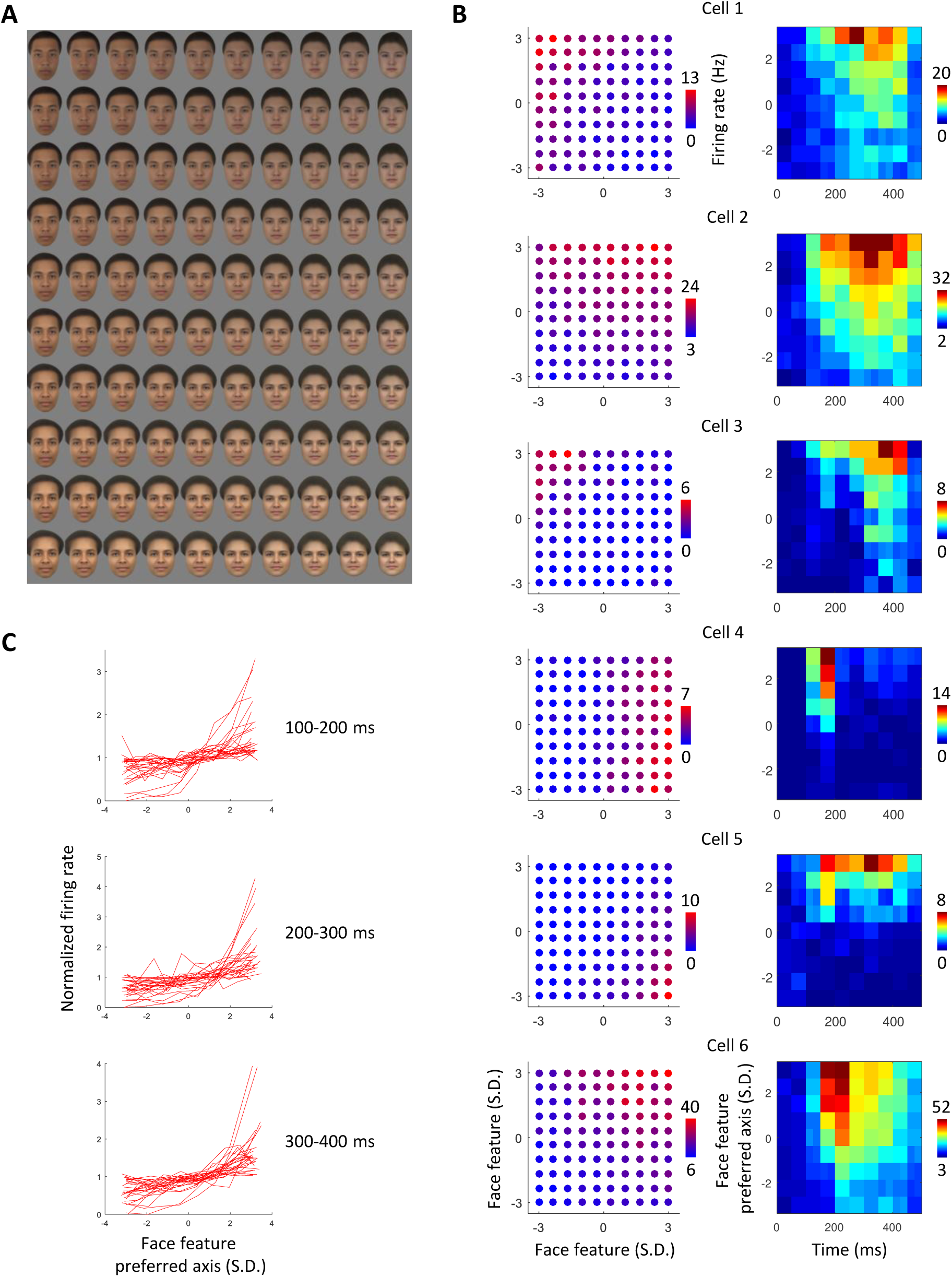
Feature tuning of AM cells measured with a longer stimulus presentation time (300 ms ON, 300 ms OFF), demonstrating robust axis tuning. **(A)** Example stimulus set consisting of 10 x 10 faces evenly spanning a 2D plane of face space ranging from - 3 SD to 3 SD. Five randomly sampled face planes were presented. **(B)** Responses of six example cells to the 2D face plane eliciting maximum modulation. (Left) Responses averaged between 50 - 300 ms. (Right) Average responses along preferred axis over time. **(C)** Response tuning curves of all cells along each cell’s preferred axis for three different time windows; firing rate was normalized by dividing by the mean firing rate.

**Figure S12.**
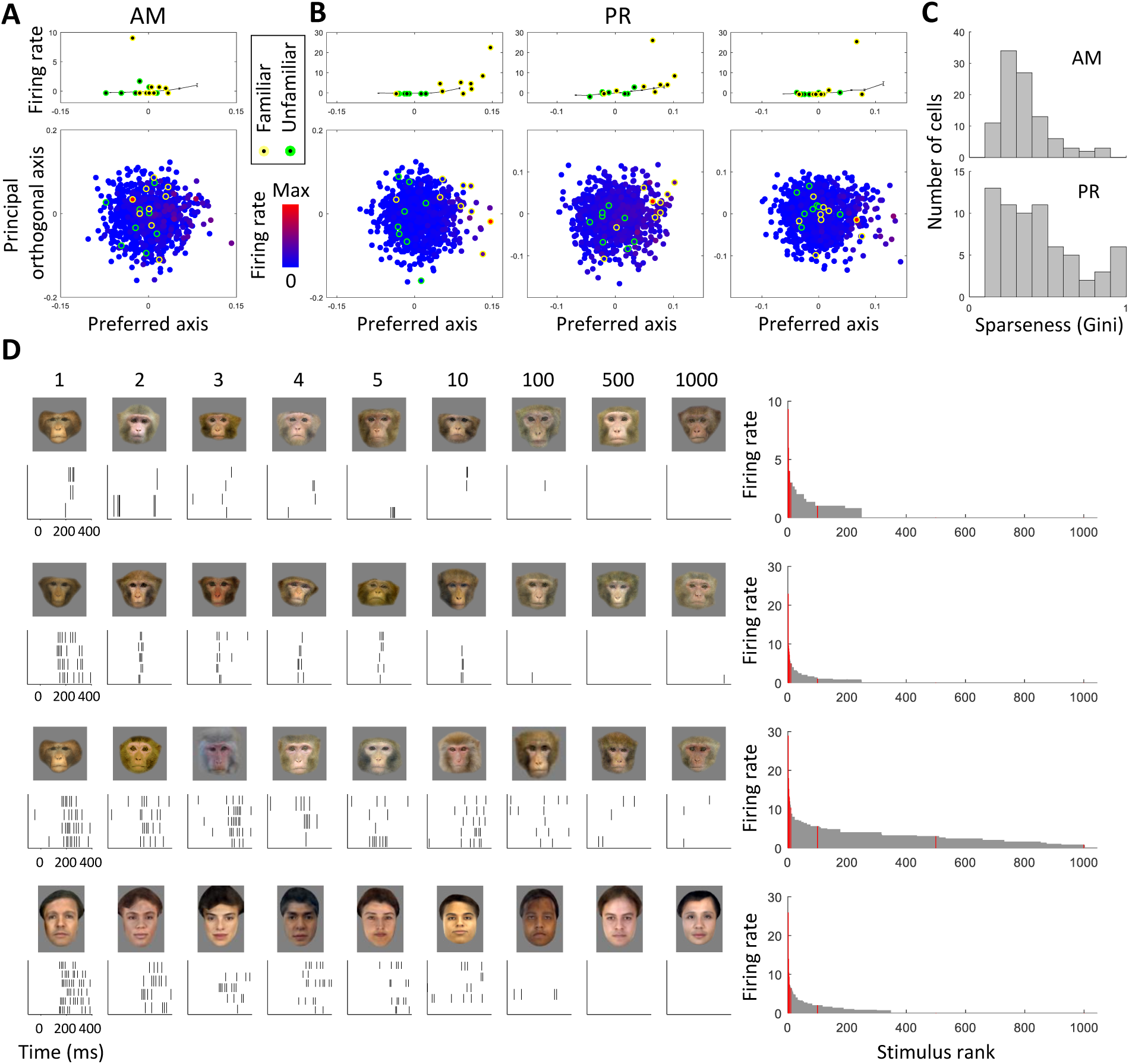
Sparsely responding AM and PR cells. **(A)** An example sparse cell from AM. **(B)** Three example sparse cells from PR. **(C)** Distributions of sparseness indices (Gini coefficient) of reliably responsive cells (reliability>0.2, see Methods). The data for this analysis included not only the 134 AM and 72 PR cells that were presented with the thousand monkey face stimulus set but an additional 53 AM and 65 PR cells that were presented with a thousand human face stimulus set (**Fig. S2B, C**) after initial screening identified them as being more responsive to human faces. We wanted to maximize our chances of identifying cells selective for specific familiar individuals, and our monkeys had personal familiarity with both humans and monkeys. **(D)** Left: Raster plots of responses of 4 example cells to 9 stimuli (five most preferred stimuli and four regularly sampled stimuli); the 4 cells from top to bottom correspond to the 4 cells in (A, B) from left to right. Right: Histograms of mean responses of each cell to the full set of 1000 stimuli; red bars indicate stimuli shown on left. [Note: the human faces in the bottom row of (D) have been replaced by synthetically generated faces due to biorxiv policy on displaying human faces.]

**Figure S13.**
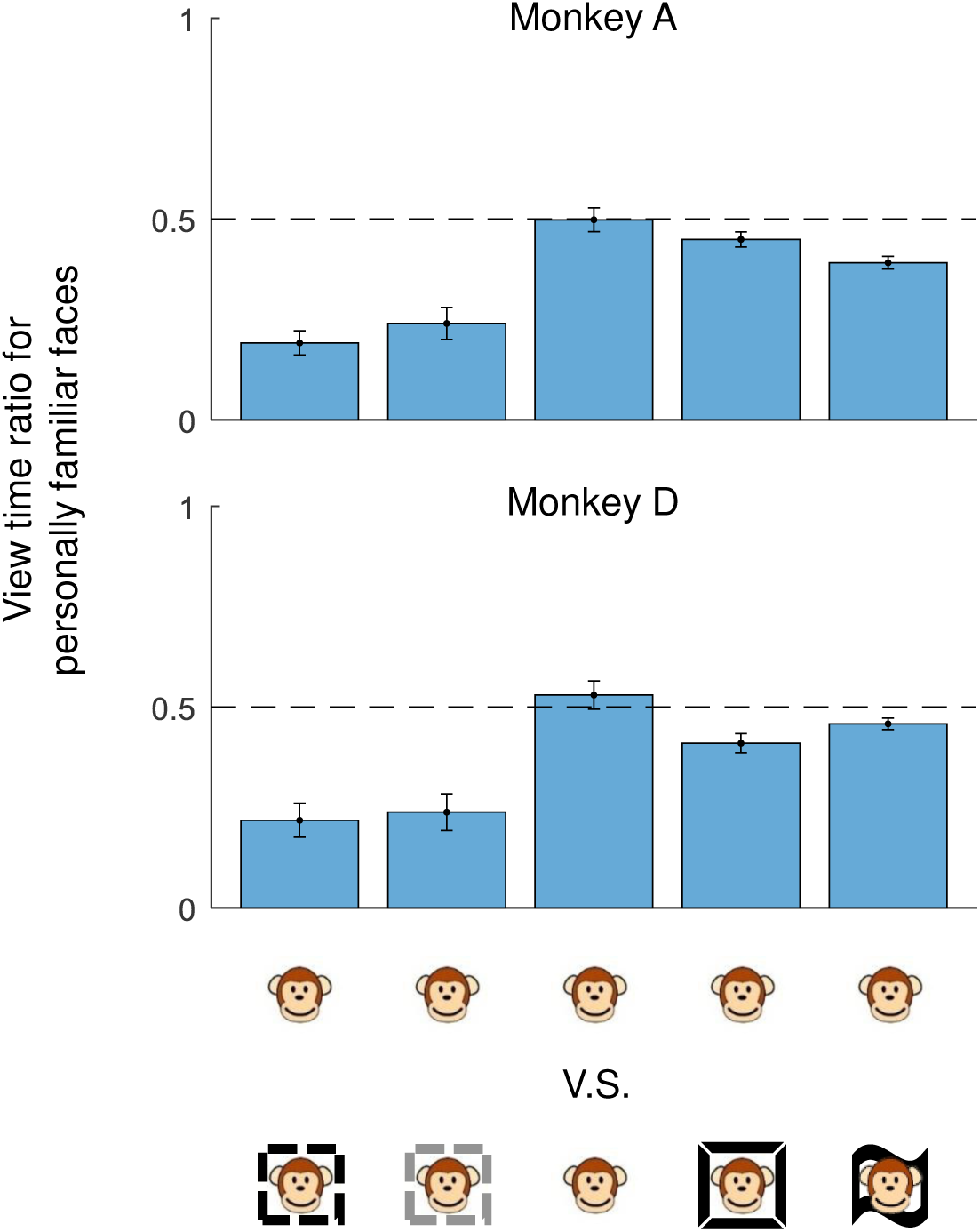
Preferential looking test. Comparing looking time to personally familiar faces versus novel unfamiliar faces, unfamiliar faces (from 1000 face set), personally familiar faces (two distinct personally familiar faces were presented on each trial), pictorially familiar faces, and cinematically familiar faces (same icon conventions as **Fig. S2**).

**Figure S14.**
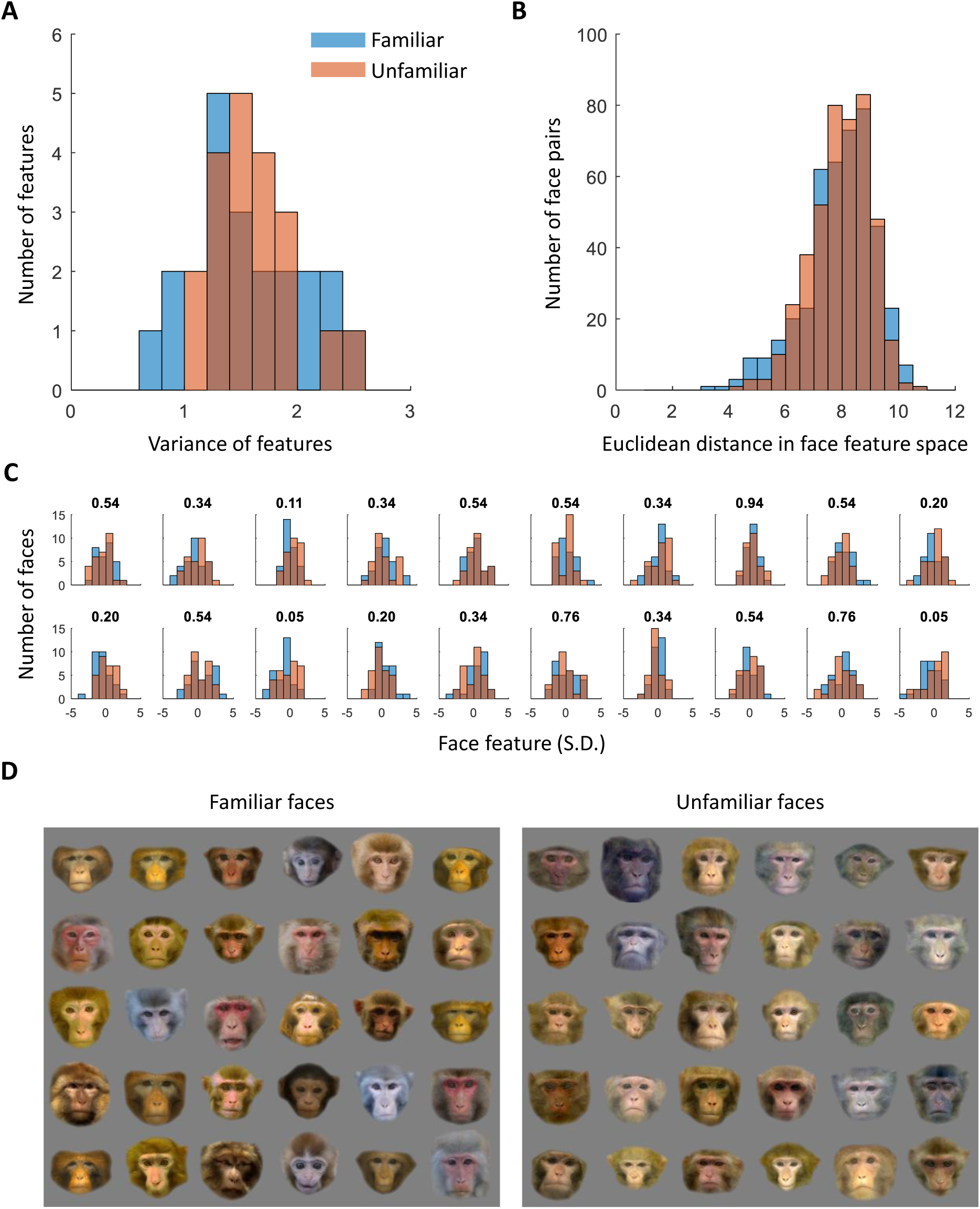
Matching the face features of familiar and unfamiliar faces. **(A)** Distribution of variances of first 20 features for 30 familiar and 30 unfamiliar feature-matched faces (K-S test, p = 0.96, N = 20). **(B)** Distribution of pairwise distances in face feature space (first 20 features) for the 30 familiar and 30 unfamiliar feature-matched faces (K-S test, p = 0.51, N = 435). **(C)** Distribution of top 20 feature values for the 30 familiar and 30 unfamiliar feature-matched faces; the number above each plot gives the p value of K-S test (N = 30) between the two feature distributions. **(D)** Images of the 30 familiar and 30 unfamiliar feature-matched faces.

**Figure S15.**
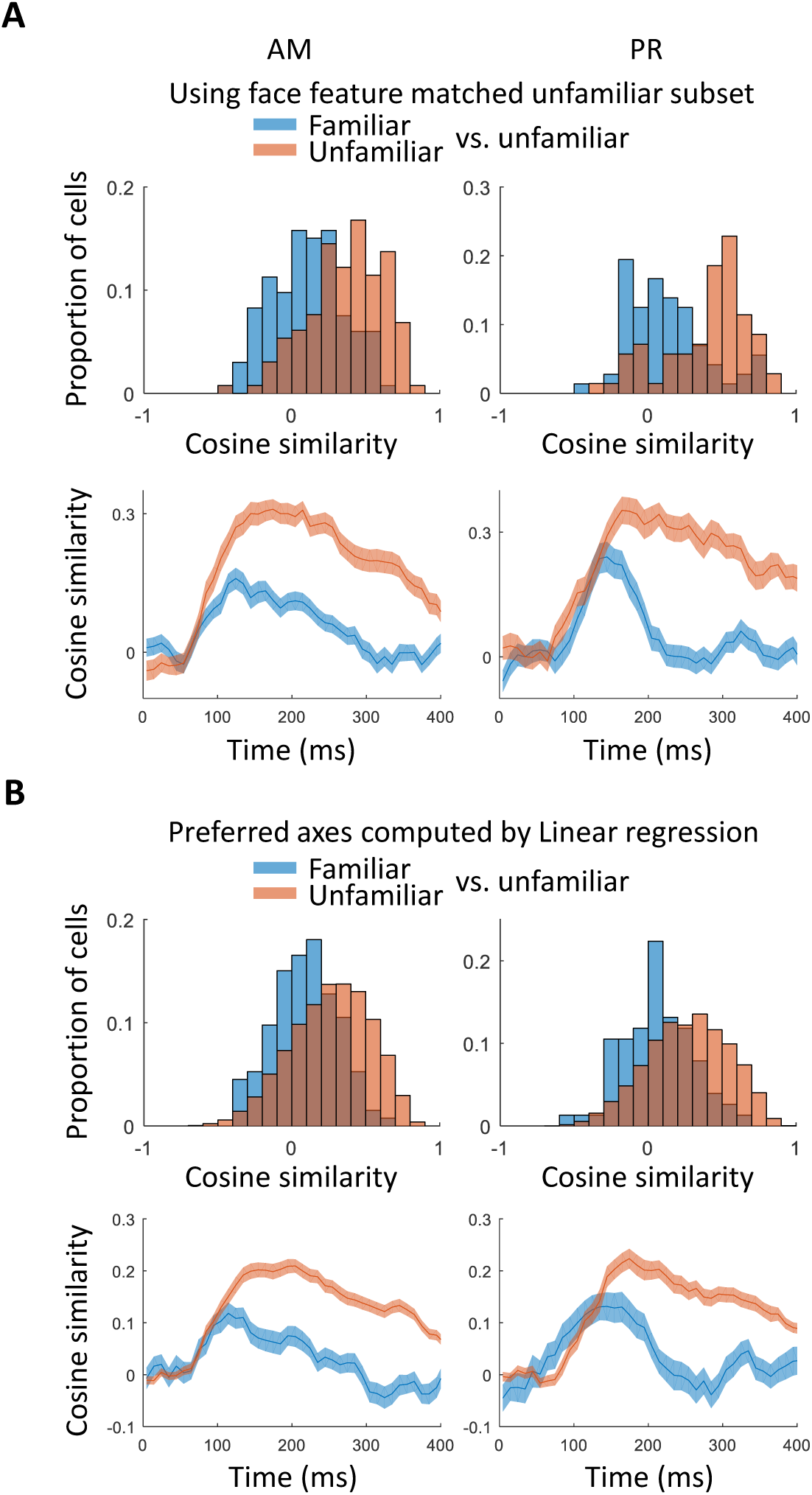
Control analyses confirming that axis change for familiar vs. unfamiliar faces was not due to stimulus differences or method of computing preferred axis. **(A)** Top: population analysis of preferred axes for familiar versus unfamiliar faces; same conventions as in Fig. 2B except 30 familiar and 30 unfamiliar feature-matched faces were used (see Methods and **Fig S14**). Bottom: time course from the same analysis; same conventions as in Fig. 2D. **(B)** Top: population analysis of preferred axes for familiar versus unfamiliar faces, same conventions as in Fig. 2B except the preferred axes were computed using linear regression rather than spike-triggered averaging (see Methods). Bottom: time course from the same analysis, same conventions as in Fig. 2D.

**Figure S16.**
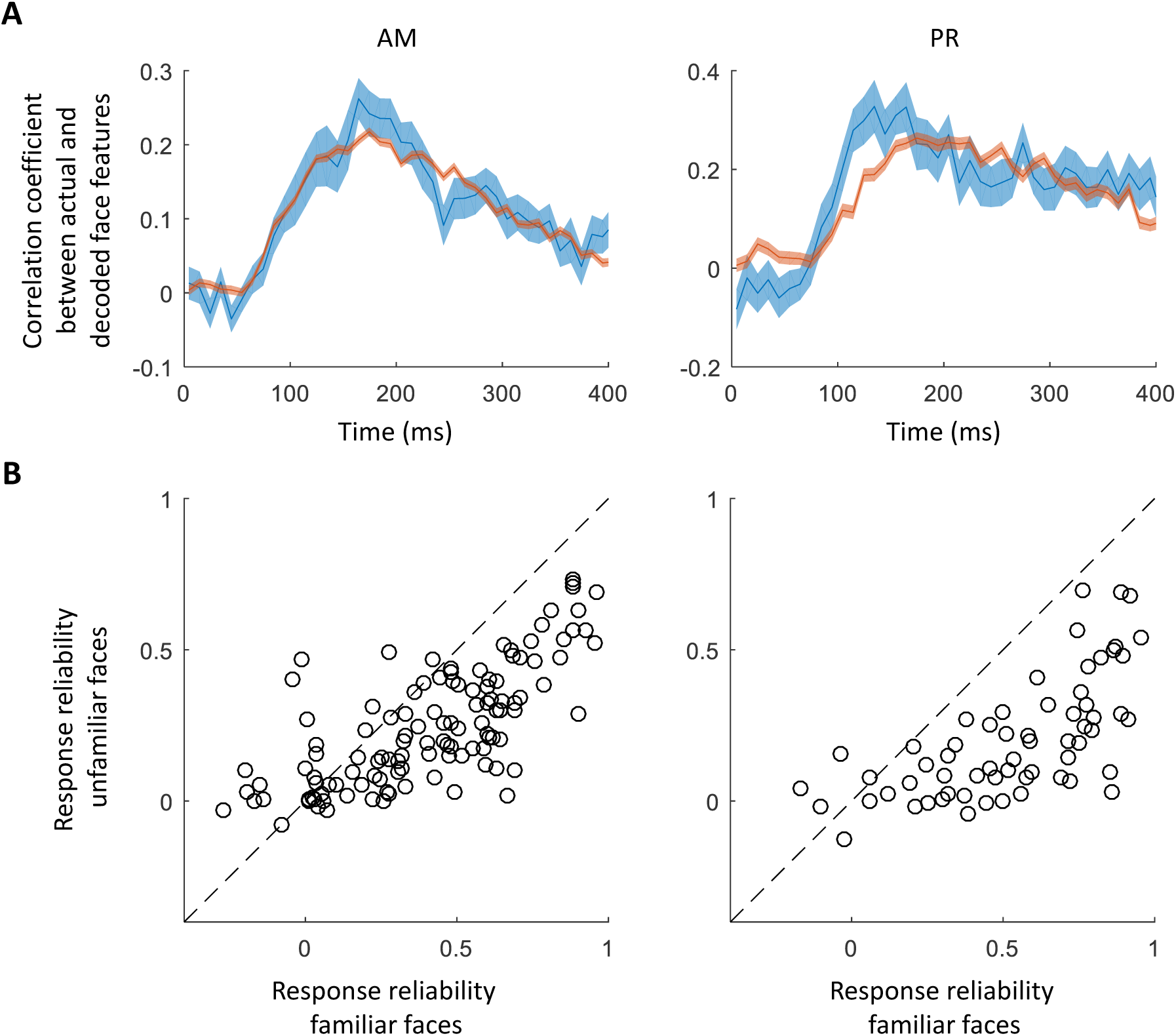
Control analyses analyzing face decoding performance and response reliability for familiar and unfamiliar faces. **(A)** Time course of linear decoding performance for familiar and unfamiliar faces measured by the correlation coefficient between actual and decoded face feature vectors, as in Fig. 2**E1**, except here, the decoder was trained using both familiar and unfamiliar faces. To match performance of familiar and unfamiliar decoding, in AM (PR), 36 (36) familiar and 150 (300) unfamiliar faces were used, sampled 10 times from the set of 1000. **(B)** Response reliability of responses to familiar versus unfamiliar faces. Each circle represents one cell. The familiar reliability was computed using 36 familiar faces, the unfamiliar one was the mean reliability of 100 repeats of 36 randomly-sampled unfamiliar faces.

**Figure S17.**
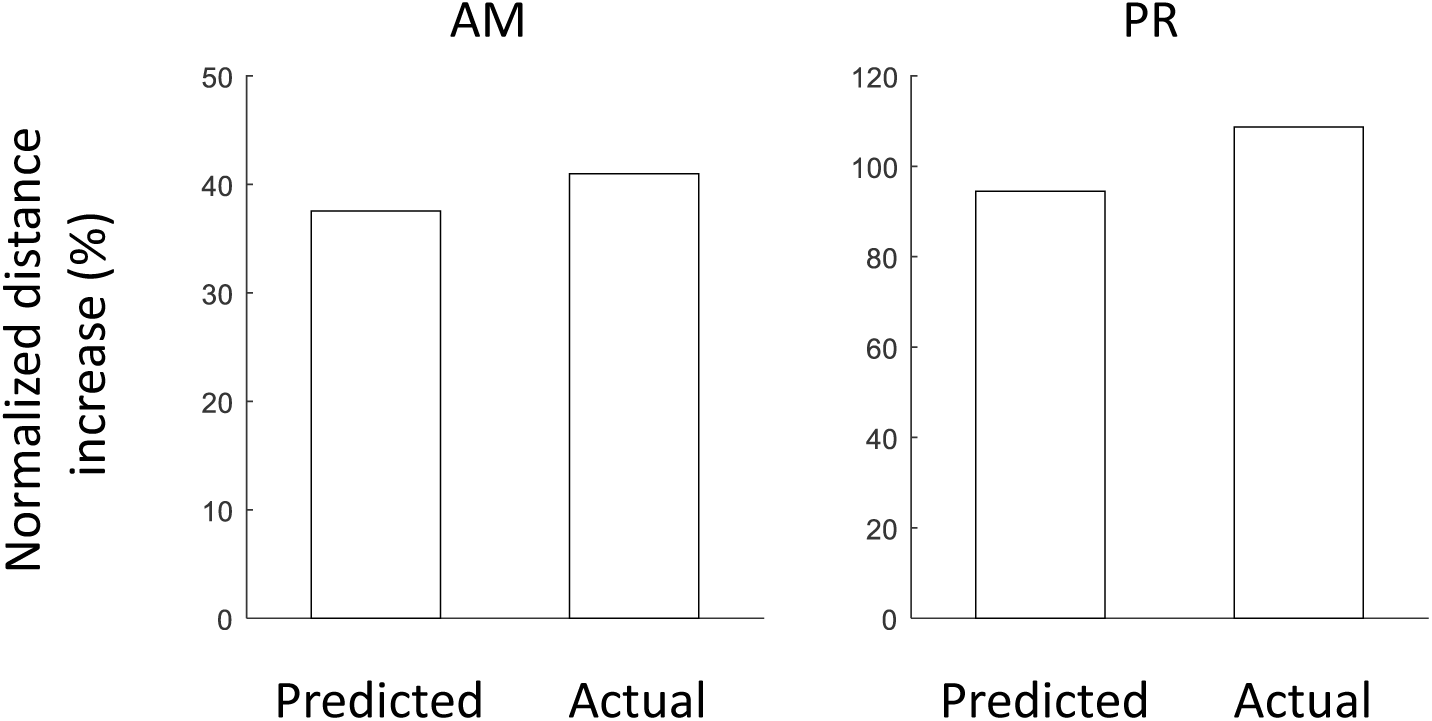
Distance increase for familiar faces using linear-model-predicted or actual response during full time window (50-300 ms). Distance increase was quantified by *(D_familiar_ – D_unfamiliar_) / (D_unfamiliar_ - D_baseline_),* where *D_baseline_* is the mean distance between unfamiliar faces during 0-50 ms.

**Figure S18.**
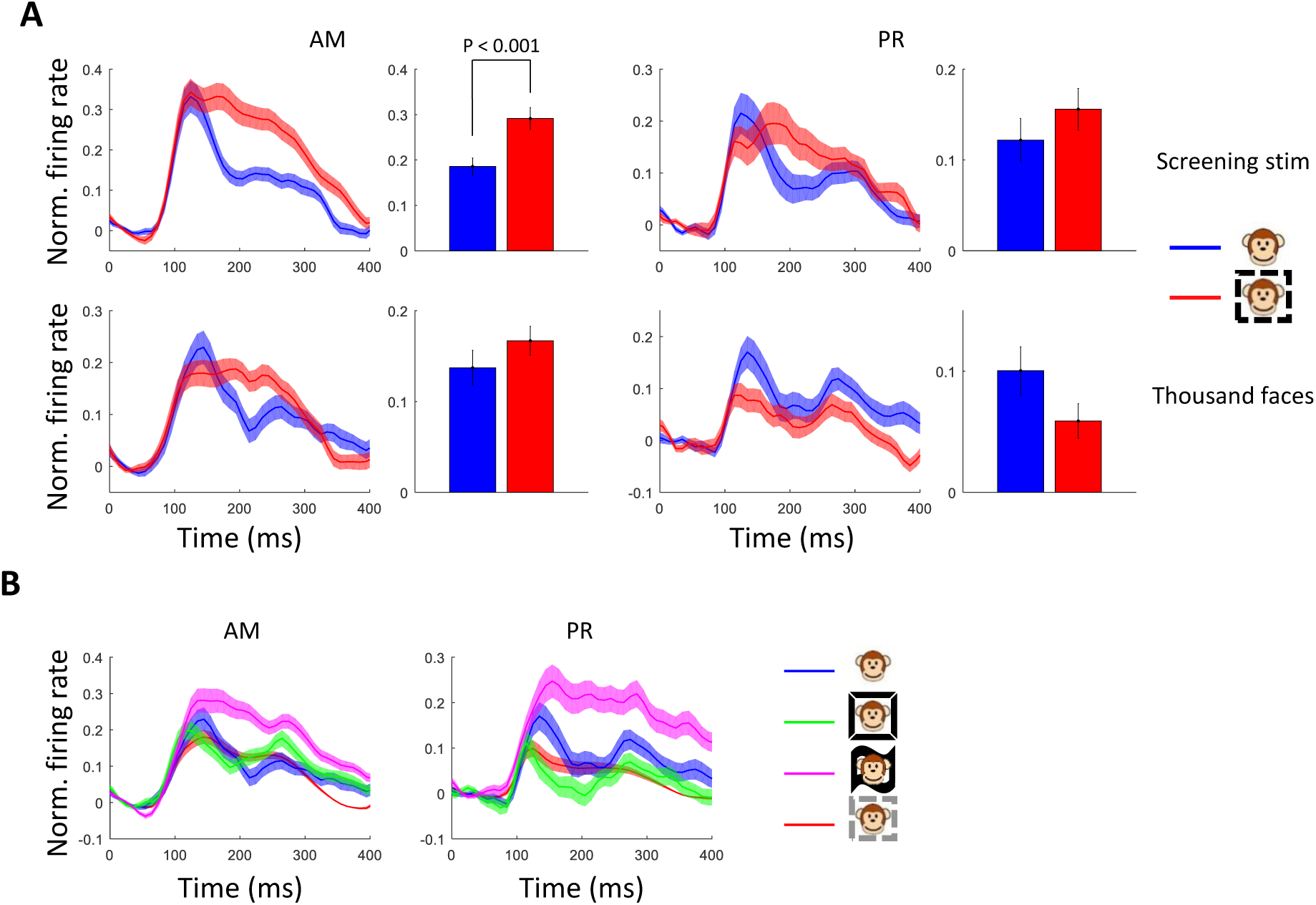
Analysis of familiar face responses separated by familiarity subtype. **(A)** Comparison of average response time courses in AM and PR to the exact same set of familiar and unfamiliar stimuli, presented in two different temporal contexts. Bar graphs: average over time window [100 300] ms. Top: Responses to 9 personally familiar and 8 unfamiliar monkey faces presented as part of screening stimulus experiment. Bottom: responses to the same set of stimuli presented as part of thousand face stimulus experiment. Shaded area, SEM**. (B)** Response time courses to faces from AM and PR populations, separated by familiarity subtype. Responses were averaged across cells and across each subtype of face from the 1000 monkey face experiment. The differences between response time courses to the three familiarity subtypes further underscores that relative magnitude of mean response to familiar faces is highly context-dependent. Shaded area, SEM.

**Figure S19.**
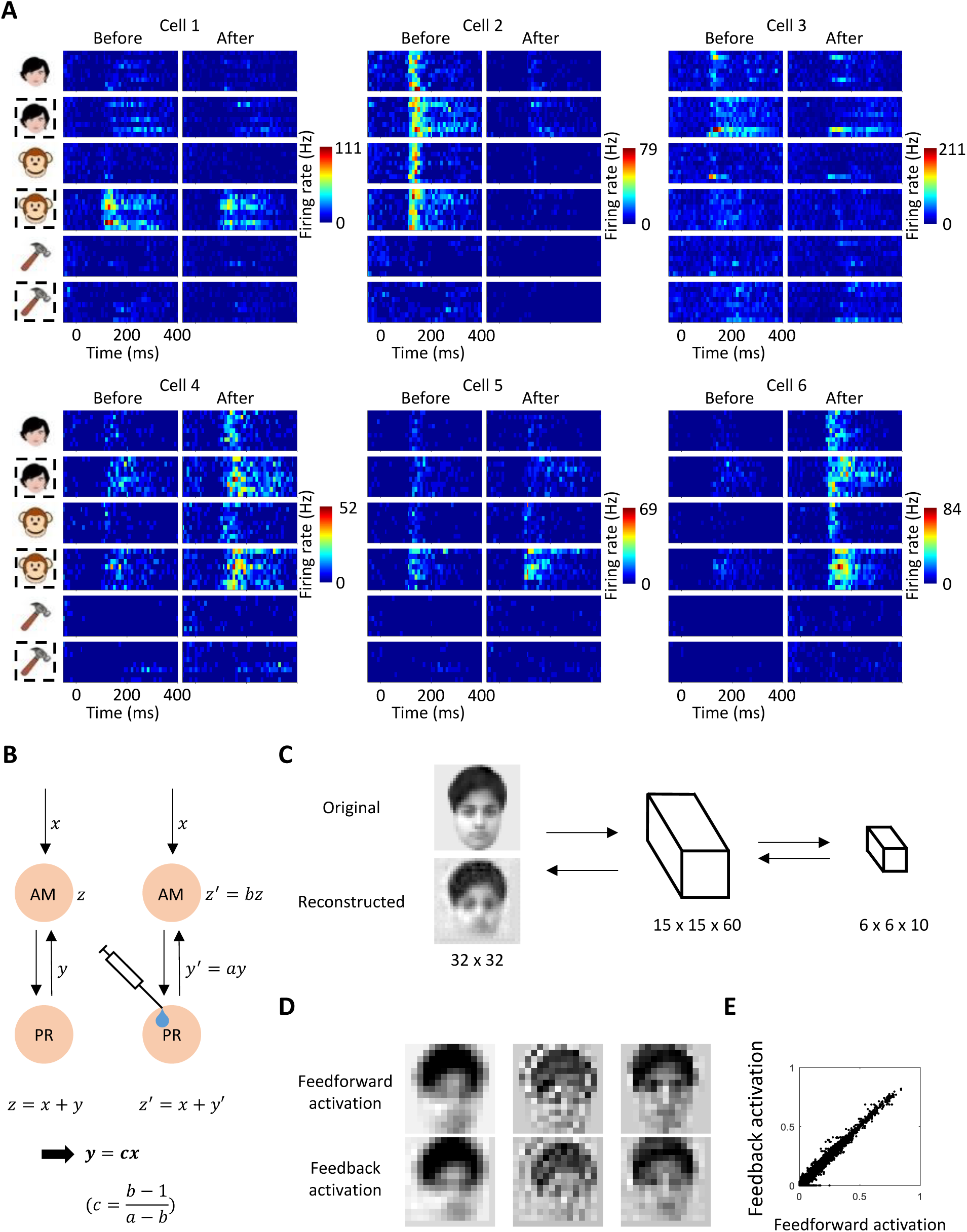
Effects of PR inactivation on AM responses are consistent with a generative model. **(A)** Responses of six additional AM example cells to screening stimuli before and after muscimol injection (same conventions as Fig. 4B). **(B)** Schematic illustration showing that feedback input from PR to AM should be proportional to the feedforward input to AM, under the empirical observation that silencing PR simply scales the response in AM (i.e., *z’ = bz*). **(C)** Schematic illustration of stacked convolutional auto-encoder *(67)*, with an example image and resulting reconstruction. The auto-encoder embeds an input image in two sequential stages, first into 60 channels of 15 x 15 features, then into 10 channels of 6 x 6 features. The process is then reversed, and the reconstructed image is generated from the embedding of the final stage. **(D)** Activation of 3 example channels in the middle stage during feedforward pass (upper row) and feedback pass (bottom row) of the auto-encoder processing. The input image is the same as in (C). Note the high similarity between the feedforward and feedback signals. **(E)** Feedforward and feedback activation of all units of the middle stage to the example image in (C). [Note: the human face in (C) is synthetic and conforms to biorxiv policy.]

**Figure S20.**
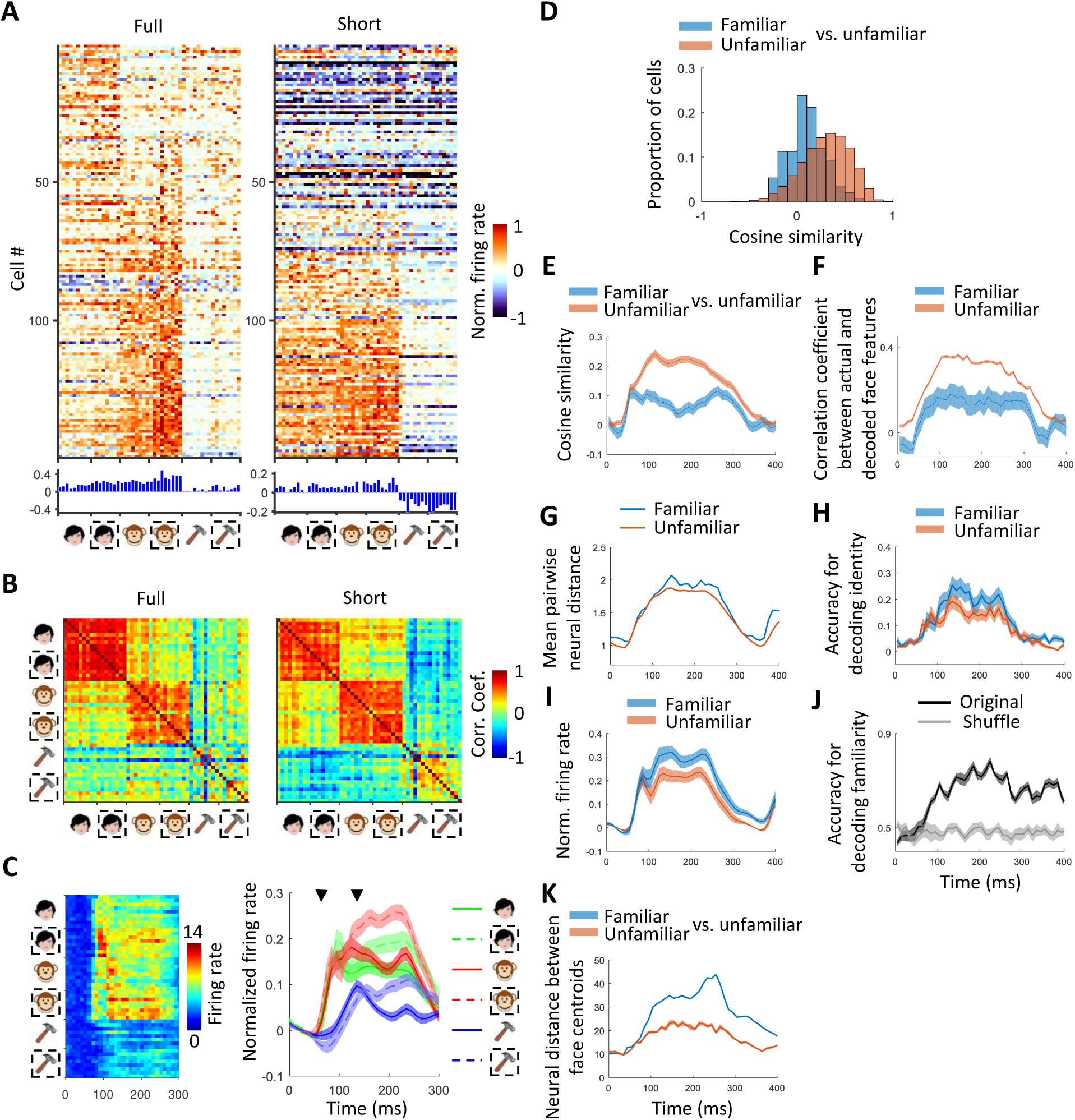
Representation of familiar stimuli in face patch ML. **(A)** Responses of cells to screening stimuli from six stimulus categories (familiar human faces, unfamiliar human faces, familiar monkey faces, unfamiliar monkey faces, familiar objects, and unfamiliar objects), recorded the face patch ML. Left, responses were averaged between 50 to 300 ms after stimulus onset (“full” response window). Right, same for a “short” window 50 to 125 ms. **(B)** Similarity matrix of population responses for full response window (left) and short response window (right). **(C)** Left: Average response time course across the ML population to each of the screening stimuli. Right: Response time course averaged across cells and category exemplars. Earlier arrow indicates the mean time when visual responses to faces became significantly higher than baseline (77.5 ms). Later arrow indicates the mean time when responses to familiar versus unfamiliar faces became significantly different (175 ms and 145 ms for human and monkey faces, respectively). Responses also diverged briefly at very short latency (95 ms and 105 ms for human and monkey faces, respectively). **(D)** Population analysis comparing preferred axes for familiar versus unfamiliar faces. Same conventions as Fig 2B. **(E-K)** Same analyses for the ML population (N = 154 cells) as in Fig. 2D**-G,** Fig. 3B**-D**.

**Figure S21.**
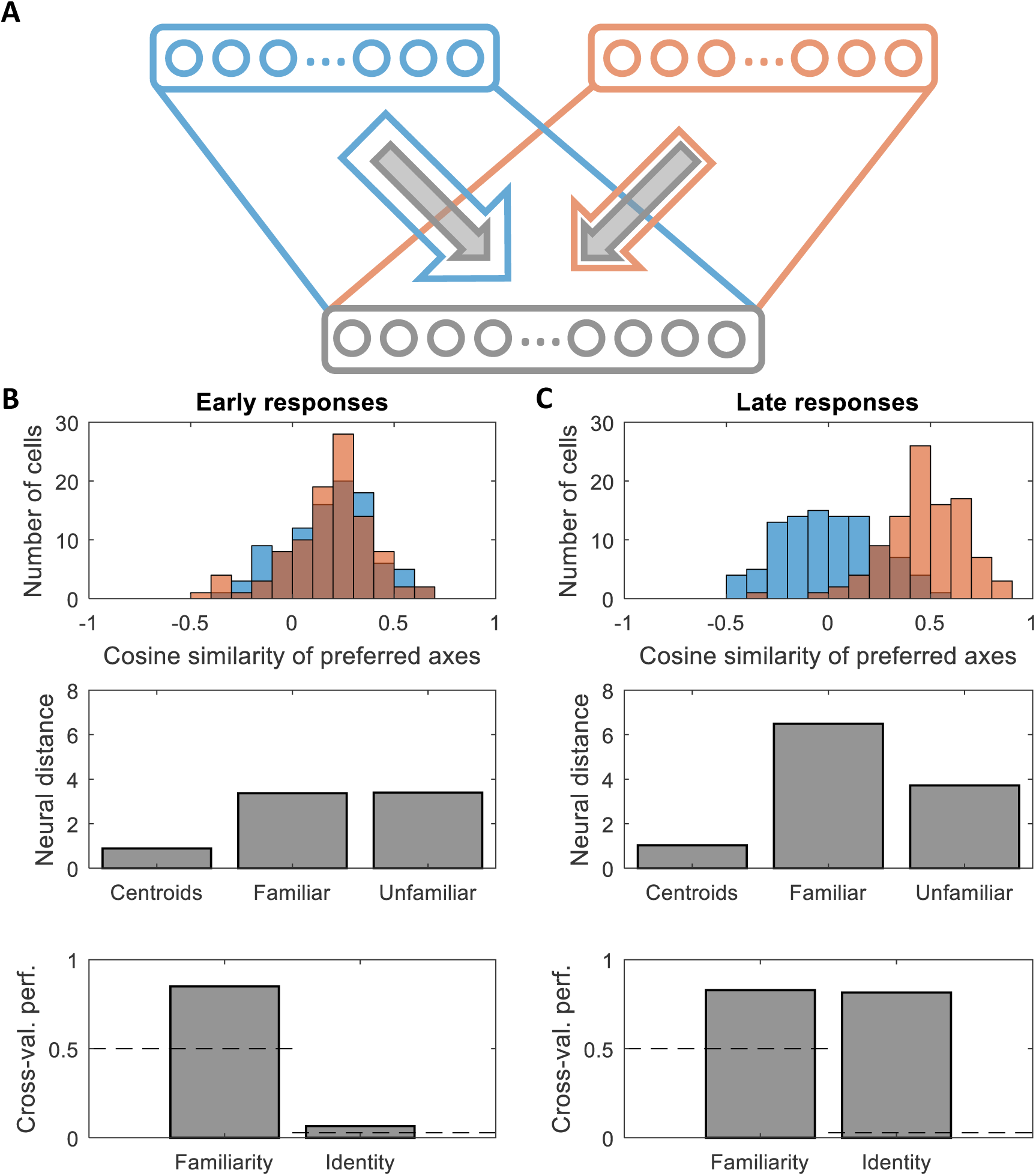
A simple network model of joint representations of face features and familiarity. **(A)** Schematic of the neural network. Populations of firing rate neurons that linearly encode face features (top) project via random (i.i.d. Gaussian) synaptic weights to the face patch neurons (bottom), which are modeled using sigmoidal activation functions. As long as the firing rates of the face patch neurons remain approximately within the linear regime of the sigmoidal non-linearity, they will inherit from their inputs a representation of the face features that lies in an approximately linear subspace of their firing rate space. **(B)** Short latency representations and their properties. The strength of the inputs from the feature coding units at short latency are independent of the familiarity of the stimulus (gray arrows in A). However, each face patch neuron also receives a feature-independent input (bias) whose sign is modulated by the familiarity of the currently presented face. This introduces a relative shift of the coding manifolds for familiar and unfamiliar faces. Because some neurons increase their firing rate with familiarity while others decrease their firing rate, this shift need not be correlated with the overall firing rate of the population. In the top panel we repeat the analysis of Fig. 2B, comparing the preferred axes in response to familiar faces to those for unfamiliar faces (blue), which yields a very similar distribution of cosine similarities as the control using a subset of the unfamiliar faces (orange). In the middle panel we show the distances in firing rate space between the centroids of the familiar and unfamiliar coding space, as well as the mean pairwise distances between two familiar face representations and two unfamiliar ones, as in Fig. 2F. These mean pairwise distance are very similar in magnitude. The bottom panel shows the decoding performance for familiarity of a linear classifier (cross-validated on faces not used for training), as well as the decoding performance of a multi-class support vector machine for the identity of a particular familiar face (out of all 36, cross-validated on held-out trials). The former is large due to the shift between familiar and unfamiliar coding spaces that enables generalization of familiarity across faces, while the latter, though above the chance level of 1/36, is relatively small (compare to Fig. 3C, Fig. 2G). **(C)** Long latency representations and their properties. Same as panel (B), except that here we model the late responses of the face patch neurons using separate input pathways from the feature-coding units (with different sets of random weights) that are active either for familiar faces (blue arrow in panel A), or for unfamiliar faces (orange arrow in A), while for panel B those pathways were activated in a familiarity-independent fashion. The relative shift of the familiar and unfamiliar coding manifolds is still present, and in addition we introduce a small constant bias of familiar face responses to model the increased average firing rate for familiar faces at long latency (see Fig. 3A**, B**). Due to the separate input pathways, the cosine similarities of the preferred axes in response to familiar and unfamiliar faces (top panel, blue) are now centered around zero, and substantially smaller than for the control (orange). The distances in firing rate space between pairs of familiar faces (middle panel) are substantially larger than for pairs of unfamiliar faces. This increased distance supports a much better decoding performance for the identity of individual familiar faces (bottom panel).

**Figure S22.**
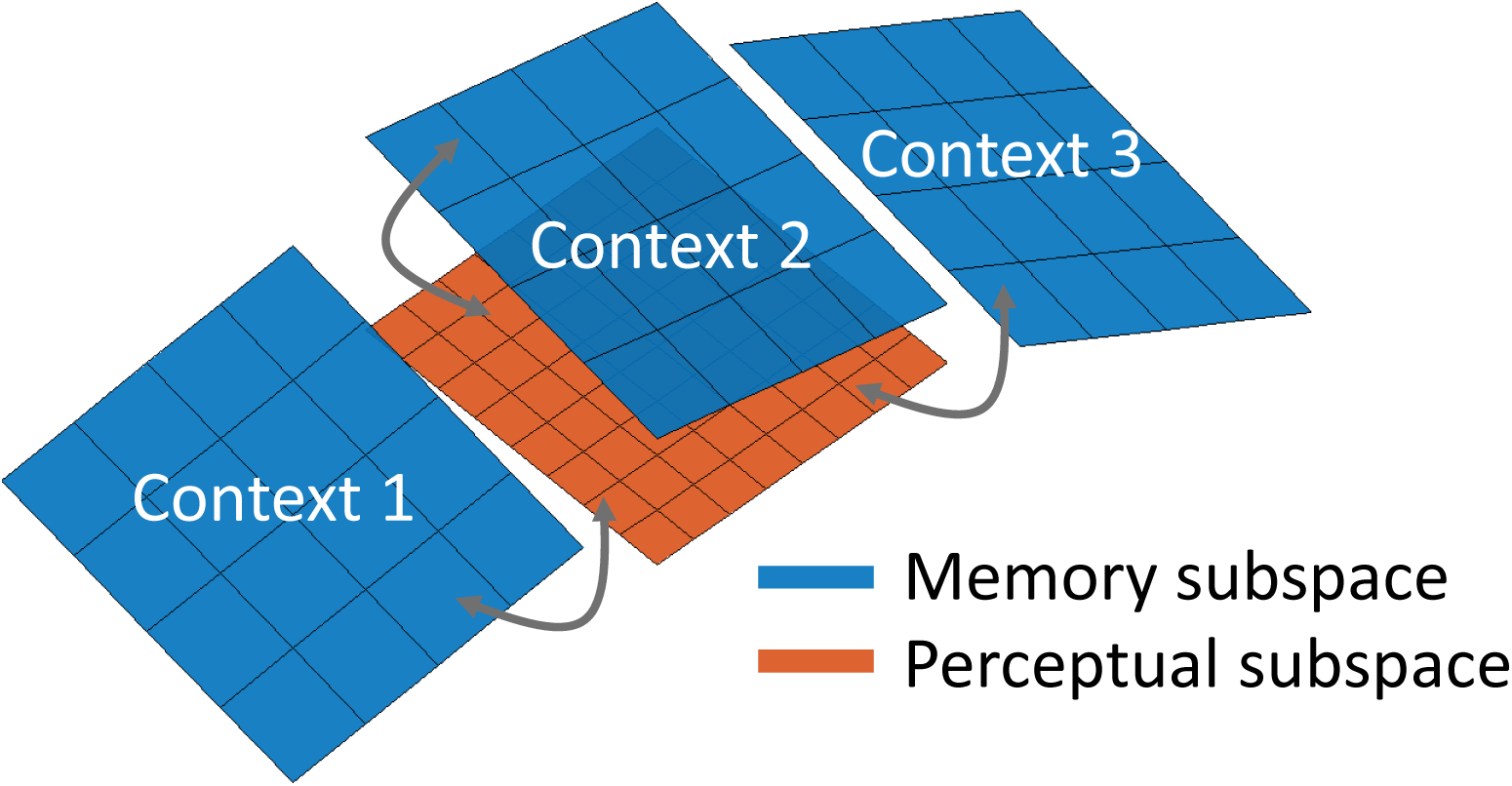
Different contexts may invoke representations in different subspaces. Each context shifted from that for representing unfamiliar stimuli. In the current study, we examined representation of familiar faces in a highly impoverished context in which the familiar stimuli were presented on a gray background during passive fixation. It is possible that under more naturalistic presentation conditions, distinct behavioral contexts (e.g., different environments) may each invoke distinct memory-gated subspaces.

**Figure S23.**
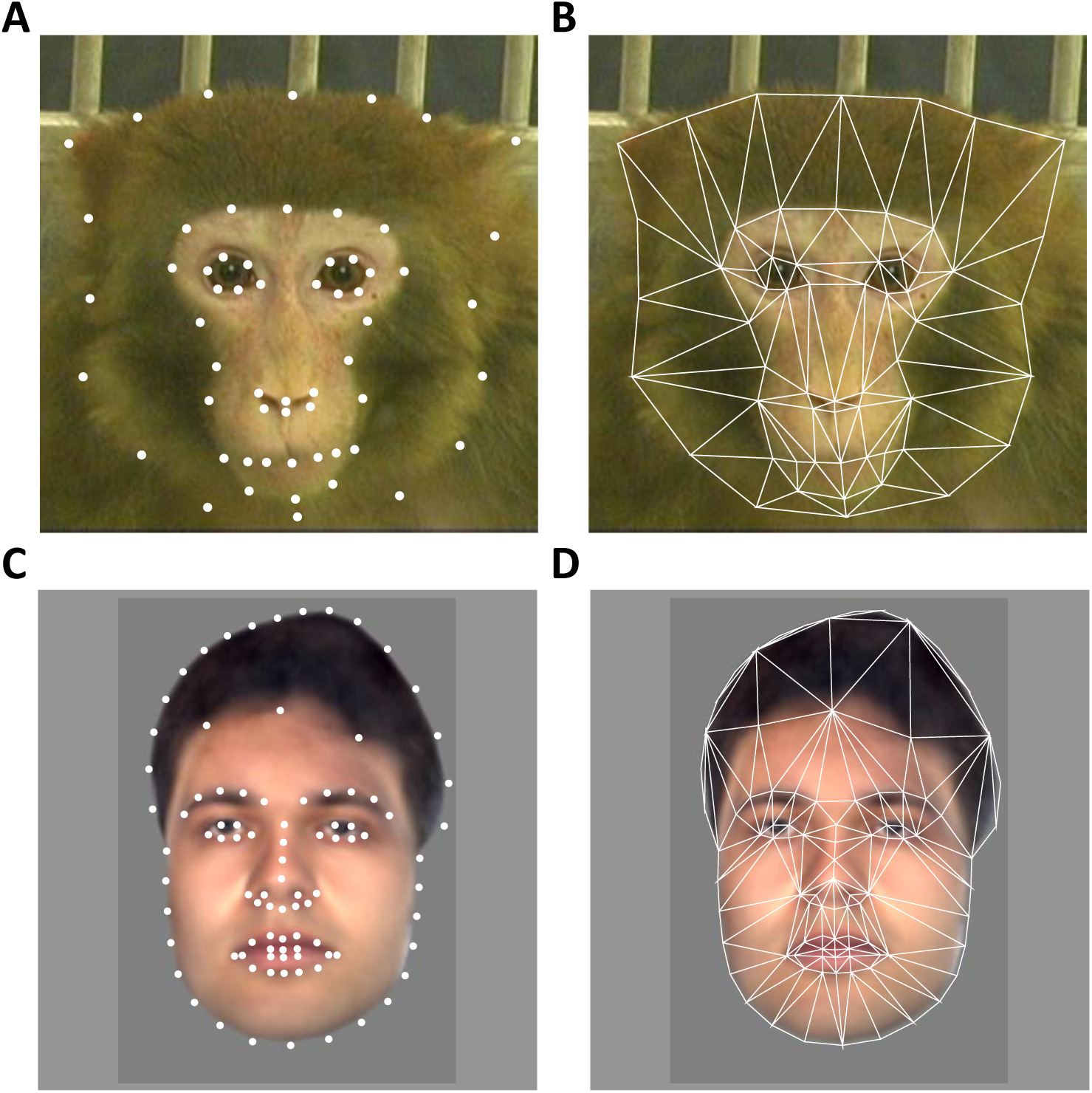
Face landmarks and shape meshes. **(A)** Example monkey face showing landmarks used as the “shape” parameters in monkey face model. **(B)** Example monkey face showing the shape mesh used for morph the face to the landmark template in the face model (see Methods). **(C)** Landmarks for an example human face. **(D)** Shape mesh for an example human face. [Note: the human face in (C, D) is synthetic to conform to biorxiv policy and was not one of the human faces actually used in the experiments.]

